# Molecular and morphological characterisation of four new ancyromonad genera and proposal for an updated taxonomy of the Ancyromonadida

**DOI:** 10.1101/2023.04.29.538795

**Authors:** Naoji Yubuki, Guifré Torruella, Luis Javier Galindo, Aaron A. Heiss, Maria Cristina Ciobanu, Takashi Shiratori, Ken-ichiro Ishida, Jazmin Blaz, Eunsoo Kim, David Moreira, Purificación López-García, Laura Eme

## Abstract

Ancyromonads are small biflagellated protists with a bean-shaped morphology. They are cosmopolitan in marine, freshwater and soil environments, where they attach to surfaces while feeding on bacteria. These poorly known grazers stand out by their uncertain phylogenetic position in the tree of eukaryotes, forming a deep-branching ‘orphan’ lineage that is considered key to better understanding the early evolution of eukaryotes. Despite their ecological and evolutionary interest, only limited knowledge exists about their true diversity. Here, we aimed to better characterise ancyromonads by integrating environmental surveys with behavioural observation and description of cell morphology, for which sample isolation and culturing is indispensable. We studied 18 ancyromonad strains, including 14 new isolates and 7 new species. Most of them belong to three new and genetically divergent genera: *Caraotamonas*, *Nyramonas*, and *Olneymonas* (encompassing 4 species). The remaining three new species belong to the already known genera *Fabomonas* and *Ancyromonas*. We also raised *Striomonas*, formerly a subgenus of *Nutomonas*, to full genus status, based on morphological and phylogenetic grounds. For all these new species, we studied their morphology under light and electron microscopy and carried out molecular phylogenetic analyses, including as well 18S rRNA gene sequences from several environmental surveys. Based on these analyses, we have updated the taxonomy of Ancyromonadida.

## INTRODUCTION

Ancyromonads are single-celled eukaryotes commonly found in diverse marine, freshwater, and soil environments (Lee and Patterson, 2000; Scheckenbach et al., 2006; Stock et al., 2009; Tikhonenkov et al., 2006). At first glance, ancyromonad cells are tiny gliding heterotrophic biflagellates (Archibald et al., 2017). Their characteristic traits are a rounded to bean-shaped cell body with a rostrum (either a distinct anterior part in a rounded cell, or the anterior lobe of a bean-shaped one) containing extrusomes, and a single prominent flagellum inserted through a fold made by the rostrum at the anterior end of the cell. The cells are strongly flattened and rigid, and have a pellicle under the cell membrane composed of a single layer of densely-staining material. Cells anchor themselves by the tip of the posterior flagellum and oscillate rapidly about the point of flagellar insertion. The distal parts of both anterior and posterior flagella are usually acronematic (thinner than the proximal part of the flagellum), and, in all investigated cases, contain only the central pair from the axoneme.

*Ancyromonas sigmoides* was originally described by Saville Kent (1882) but reported only sporadically thereafter. The modern concept of the genus was established almost a century later, by Hänel in 1979 (Hänel, 1979; translated from German in Heiss et al., 2010). Organisms fitting this description have been found in widely varying locations around the world (Lee and Patterson, 2000; Prokina and Philippov, 2018; Scheckenbach et al., 2006; Simpson and Patterson, 2006). However, Cavalier-Smith et al. (2008) argued that the organism identified by Hänel as *A. sigmoides* was morphologically distinct from, and probably unrelated to, the organism in Kent’s original description. This was disputed, resulting in both a reassessment of the taxonomy of *Ancyromonas* and the recognition of greater diversity within the group than had previously been appreciated (Cavalier-Smith et al., 2008; Glücksman et al., 2013; Heiss et al. 2010, 2016, Torruella et al. 2017). At present, the ancyromonad strain B-70 has been described as the neotype of *A. sigmoides* (Heiss et al., 2010). Other ancyromonads have been placed in the genera *Planomonas, Fabomonas,* and *Nutomonas* (Glücksman et al., 2013), the latter containing all known ancyromonads from freshwater environments.

Despite its cosmopolitan distribution, the true genetic and morphological diversity encompassed in the ancyromonad clade remains largely unknown. In addition, its precise position in the tree of eukaryotes is still enigmatic and debated (Atkins et al., 2000; Cavalier-Smith et al., 2014; Paps et al., 2013; Torruella et al. 2015). Ancyromonads do not belong to any of the well-accepted “supergroups”, and the most recent phylogenomic analyses suggest that they form one of the earliest-diverging branches of the tree (Brown et al., 2018), rendering them key to tackling deep evolutionary questions. Some of the difficulty in resolving the phylogenetic position of ancyromonads stems from the limited molecular data available for this lineage. This is due, in large part, to the challenge in identifying, isolating and culturing these organisms from natural samples.

Here, we report 14 new isolates of ancyromonads from marine and freshwater habitats around the world. We characterise these morphologically and molecularly, and place them into a phylogenetic framework containing all ancyromonad diversity known from ribosomal RNA environmental surveys.

## MATERIALS AND METHODS

### Sampling, isolation, and culture conditions

Fourteen strains of ancyromonads were collected from sediment, freshwater and marine water samples. A small amount of each environmental sample was inoculated into 1% YT medium (100 mg yeast extract, 200 mg tryptone, 100 ml distilled water) or Cerophyl^®^ medium (ATCC 802) made from distilled water or natural seawater (corresponding to the sample source) and cultured at room temperature. Some strains were enriched through as many as five rounds of 1:5 serial dilution in 1% YT medium. Individual ancyromonad cells were isolated from enrichment cultures either manually with a micropipette or with an Eppendorf PatchManNP2 micromanipulator using a 65-µm VacuTip microcapillary (Eppendorf) under a Leica Dlll3000 B inverted microscope, and inoculated in 96-well plates.

Cultures of the new isolates were maintained alongside four previously characterised cultures (*Ancyromonas sigmoides* B-70, *Ancyromonas kenti* EDM11b, *Fabomonas tropica* NYK3C, and *Striomonas longa* ncfw stat. nov. [formerly in *Nutomonas*]) (Table 1). All marine isolates except for *A. sigmoides* B-70 were maintained in sterile seawater with 1% YT medium and co-cultured prokaryote prey from corresponding source samples. The *A. sigmoides* strain B-70 was grown in a mixture of sterile seawater and Volvic^®^ mineral water (1:1) containing 1% YT medium. All marine strains were subcultured every week. Freshwater strains were maintained in Volvic^®^ mineral water with 1% soil extract and subcultured every two to three days. All cultures were maintained in 50-ml culture flasks at 15°C. All new strains described in this work are available upon request from the culture collection of the DEEM Team, University of Paris-Saclay, France.

**Table 1.** Isolates and sampling sites for all cultured ancyromonad strains characterised in this study.

| Taxonomic name | Isolate | Culture collection | Sampling site | Sampling environment | Accession number |
| --- | --- | --- | --- | --- | --- |
| *Ancyromonas sigmoides* | B-70 * | CCAP 1958/3 | near Srednii Island, Karelia, Russia | Littoral waters | MW872731 |
| *Ancyromonas kenti* | EDM11b * | CCAP 1953/1 | Burrow-in-Furness, Cumbria, England | Sediment | JQ340339 |
| *Ancyromonas* sp. | JF-An-6 | DEEM | ~ beach in New Jersey, USA | Sediment | OP740703 |
| *Ancyromonas kenti* | AKROSKO2018 | DEEM | Roc'h ar Bleiz, Roscoff, France | Intertidal sediment | MW872730 |
| *Ancyromonas mediterranea* sp. nov. | C362 | DEEM | Villefranche Bay, France | Mesopelagic water column (250m depth) | MW872728 |
| *Ancyromonas pacifica* sp. nov. | SRT-303 | DEEM | Lagoon on Iriomote Island, Okinawa, Japan | Muddy pool | OP740701 |
| *Caraotamonas croatica* gen. et sp. nov. | CRO19S5 | DEEM | Malo jezero, Mljet island, Croatia | Sediment of saltwater lake | MW872725 |
| *Nutomonas limna* | ORSAYFEB19A NCY | DEEM | University campus, Orsay, France | Ditch water | MW872726 |
| *Striomonas longa* stat. nov. | ncfw * | CCAP1958/5 | Boiling Springs, North Carolina, USA | Sediment | MW872732 |
| *Nyramonas glaciarium* gen. et sp. nov. | ORSLAND19S1 | DEEM | Solheimajokull glacier, Iceland | Sediment from melting glacier | OP740706 |
| *Nyramonas silfraensis* sp. nov. | ORSLAND19S2 | DEEM | Silfra rift, Iceland | Sediment from the walls of the rift (~2m depth) | MW872729 |
| *Olneymonas thurstoni* gen. et sp. nov. | STRT | DEEM | Olney Pond, Lincoln, Rhode Island, USA | Surface of rainbow trout | OP740702 |
| *Planomonas micra* | PMROSKO2018 | DEEM | Roc'h ar Bleiz, Roscoff, France | Sediment in a tide pool | MW872727 |
| *Planomonas* sp. | EK-PEI | DEEM | ~ beach, Prince Edward Island, Canada | Beach sand | OP740707 |
| *Planomonas* sp. | SRT-307 | DEEM | Lagoon on Iriomote Island, Okinawa, Japan | Surface of barracuda | OP740704 |
| *Fabomonas tropica* | NYK3C * | CCAP 1958/4 | Ankobra Beach, Ghana | Sediment | MW872733 |
| *Fabomonas* sp. | SRT-009 | DEEM | Japan | Marine | OP740705 |
| *Fabomonas mesopelagica* sp. nov. | A153 | DEEM | Villefranche Bay, France | Mesopelagic water column (250m depth) | MW872724 |
| Previously characterized strains are marked with an asterisk (*). Morphological data of B-70 is taken from Cavalier-Smith et al. (2008) and those for EDM11b, ncfw and NYK3C are taken from Glücksman et al. (2013). |  |  |  |  |  |

### Light and electron microscopy

Light microscopy (LM) was performed using a Zeiss Axioplan 2 imaging microscope equipped with a Sony α9 digital camera. Cell size was measured on 32 different cells for each strain. Calculations of cell size averages omitted the longest and the shortest cells. The acronematic part was excluded from the measurement of the posterior flagellum.

For transmission electron microscopy (TEM), cell suspensions of *Caraotamonas croatica* Cro19S5 gen. et sp. nov., *Nyramonas silfraensis* OrsLandS2 gen. et sp. nov. and *Fabomonas mesopelagica* A153 sp. nov., were mixed with 2.5% glutaraldehyde and 2% paraformaldehyde (final concentration) in 0.2 M sodium cacodylate buffer (SCB) (pH 7.2) at room temperature for 1 h. Cells were collected by centrifugation at 1,000 × *g* for 5 min and rinsed with 0.2 M SCB. The specimens were then post-fixed in 1% osmium tetroxide (final concentration) in 0.2 M SCB at room temperature for 1 h, followed by dehydration through an ethanol series. The specimens were embedded in Spurr’s resin (Agar Scientific, Essex UK). Ultra-thin sections were cut on a Leica EM UC6 ultramicrotome and double-stained with uranyl acetate and lead citrate. Ultra-thin sections were observed using a JEOL 1400 electron microscope at the Imagerie-Gif facility (Gif-sur-Yvette, France).

For scanning electron microscopy (SEM), cells of *Nyramonas silfraensis* OrsLandS2 were mixed with 2.5% glutaraldehyde (final concentraiton) in 0.2 M SCB. Specimens were mounted on circular glass coverslips coated with poly-L-lysine for 2 h on ice. The glass coverslips were rinsed with 0.2 M SCB and fixed with 1% osmium tetroxide (final concentration) in 0.2 M SCB at room temperature for 1 h, followed by dehydration through an ethanol series and drying with CO_2_ using a Leica CPD300 critical point dryer. Samples were then coated with platinum using a Leica ACE600 sputter coater, and observed with a Zeiss GeminiSEM 500 field emission SEM at the Imaging Facility of the Institut de Biologie Paris Seine (Paris, France).

### Ribosomal RNA sequence data

Ancyromonad ribosomal RNA (rRNA) gene sequence data were acquired through diverse approaches. The 18S rRNA gene was obtained by PCR amplification directly from lysates of each of the 14 novel cultured ancyromonads. For some cultured strains, the full ribosomal gene cluster was also obtained from preliminary transcriptomic and genomic data. Finally, 18S rRNA gene sequences were amplified from various environmental DNA samples. We give details about each of these approaches below.

For PCR amplification, actively growing cultures of strains JF-An-6, SRT303, OrsLand19S1, STRT, EK-PEI, SRT307, and SRT009 were concentrated by centrifugation for 15 min at 13,400 × *g* at room temperature. Cells were then lysed by heating in a microwave twice for 10 s. After removing the supernatant, the remaining 4 µl were used for nested PCR. The initial amplification was carried out with primers 82F (5’-GAA ACT GCG AAT GGC TC-3’) and 1520R (5’-CYG CAG GTT CAC CTA C-3’); the amplicons were then used as template for a second amplification reaction using primers 612F (5’-GCA GTT AAA AAG CTC GTA GT-3’) and 1498R (5’-CAC CTA CGG AAA CCT TGT TA-3’). All PCR reactions consisted of an initial denaturing period (95°C for 3 min), 35 cycles of denaturing (93°C for 45 s), annealing (five cycles at 45°C and 30 cycles at 55°C, each for 45 s), and extension (72°C for 2 min), and a final extension period (72°C for 5 min). Amplicons were sequenced through the Sanger approach (Genewiz, Leibniz, Germany), and the resulting chromatograms were analysed with Geneious 6.0.5 (Kearse et al. 2012). The seven partial sequences obtained with this procedure were deposited in GenBank under accession codes OP740701–OP740707 (see Table 1).

The full ribosomal RNA operon (comprising 18S, ITS1, 5.8S, ITS2, and 28S sequences) was obtained from preliminary transcriptomic sequence data from strains NYK3C, AKRosko2018, C362, Cro19S5, OrsayFeb19Ancy, OrsLand19S2, PMRosko2018, and A153, and preliminary genome assemblies of strains ncfw and B-70, (Torruella et al., unpubl. data; Blaz et al., unpubl. data). Operons were identified using BLASTn v2.2.26+ (Altschul et al., 1990), with the *Ancyromonas sigmoides* B-70 18S ribosomal RNA gene sequence (EU349231) as a query. Ten operon sequences obtained with this procedure were deposited in GenBank under accession codes MW872724-MW872733 (Table S3).

Finally, specific primers were designed for four ancyromonad genera: *Planomonas*, *Fabomonas*, *Ancyromonas*, and *Nutomonas* (Table S3). These primers were used to obtain partial ancyromonad 18S rRNA gene sequences from the DEEM laboratory collection of environmental DNA obtained from diverse field trips (see Torruella et al., 2017 for details). PCR amplifications of the 18S rRNA gene were performed using HotStart Taq-Platinum polymerase (Invitrogen, Carlsbad, CA, USA). PCR conditions consisted of 45 cycles of 94°C for 15 s, 55°C for 30 s, and 72°C for 2 min, with a final extension step of 72°C for 7 min. Amplicons were cloned using the TOPO TA cloning kit (Invitrogen) according to the manufacturer’s instructions. 24–48 clones per sample were chosen randomly, and the cloned fragments were amplified with vector primers and sequenced with the same forward and reverse primers initially used for DNA amplification. Amplicons were sequenced by the Sanger method (Genewiz, Leibniz, Germany). Chromatograms were analysed with Geneious 6.0.5 (Kearse et al., 2012). Poor quality end regions were trimmed using an error probability limit of 0.01%; forward and reverse sequences were assembled with the Geneious alignment tool, and vector sequence portions were eliminated using VecScreen (www.ncbi.nlm.nih.gov/tools/vecscreen/). Nine partial 18S rRNA gene sequences (789–1,463 bp) obtained with this procedure have been deposited in GenBank under accession codes MW812385-MW812393 (Table S3).

Additional data were obtained from the PR2, and SILVA (Clark et al., 2016; Guillou et al., 2013; Quast et al., 2013), and GenBank non-redundant (‘nr’) databases were mined using BLAST with default parameters and four representative ancyromonad sequences as queries (*Ancyromonas sigmoides* B-70: EU349231, *Planomonas micra* ATCC 50267: EF455780, *Striomonas longa* ncfw: JQ340331, and *Fabomonas tropica* nyk4: JQ340336). In order to identify and remove redundant and non-ancyromonad sequences, a multiple sequence alignment, also containing 28 outgroup sequences (representing putatively early-branching eukaryotes), was constructed from these data and used to obtain a phylogenetic tree with the IQ-TREE web server under the GTR+G4 model (data not shown). We identified 207 sequences: 29 from previously characterised ancyromonads and 178 from environmental sources (Table S2). Similarly, ancyromonad sequences were mined from five metabarcoding studies based on the v4 region of the 18S rRNA gene, including three studies from freshwater (David et al., 2021a, 2021b; Iniesto et al., 2021), and two from marine environments (Logares et al., 2020; Massana et al., 2015), resulting in 146 OTUs (Table S2).

### Phylogenetic analyses

We used the MAFFT web server (Katoh et al., 2019) to align a 50-taxon dataset containing 18S rRNA gene sequences: nine full and seven partial sequences obtained in this study, 29 previously characterised ancyromonads, and five outgroup sequences (*Mantamonas*, *Malawimonas*, *Diphylleia*, *Rigifila*, and *Micronuclearia*). Ambiguously aligned regions were trimmed using trimAl v1.4 with the automated1 setting (Capella-Gutiérrez et al., 2009), resulting in 1,592 sites. Phylogenetic analysis was performed using the IQ-TREE web server under the GTR+GAMMA4 model (Trifinopoulos et al. 2016); statistical support at branches was estimated using 1,000 ultrafast bootstrap replicates (UFBS). Since UFBS are known to overestimate branch support, the best-fitting model (TN+F+R3, according to the Bayesian Information Criterion) was also used to infer a phylogeny with 1,000 non-parametric bootstrap replicates (NPBS). A percent-identity matrix was also computed for the 45 ancyromonad sequences in this alignment using the Clustal Omega server (McWilliam et al., 2013) (Table S1).

We also carried out a phylogenetic analysis that included the environmental ancyromonad 18S rRNA gene sequences obtained from the data-mining surveys described above (Table S2). The combination of these with experimentally obtained data resulted in a total of 383 sequences after removal of redundancies, which were aligned using MAFFT. The alignment was manually inspected with AliView (Larsson 2014) to remove ambiguously aligned regions, resulting in 1,850 sites. Phylogenetic trees were inferred using IQ-TREE under the TIM3eR4 model with 1,000 NPBS, and under the GTR+GAMMA4 model with 1,000 UFBS.

All data files are available at figshare (10.6084/m9.figshare.22078451). Sequences have been deposited in GenBank (accession numbers given in Tables 1 and S3).

## RESULTS

Starting from aquatic samples from around the world, we isolated ancyromonad cells to establish mono-eukaryotic cultures. These comprise 14 new strains, which we examined along with four previously described strains (*Ancyromonas sigmoides* B-70, *Ancyromonas kenti* EDM11b, *Striomonas longa* ncfw stat. nov., and *Fabomonas tropica* NYK3C) (Table 1). All strains, except for those belonging to the genera *Nutomonas, Striomonas* stat. nov., *Nyramonas* gen. nov., and *Olneymonas* gen. nov., were isolated from marine environments.

### Light microscopy

We examined the morphology of all 18 strains by light microscopy (LM). Observations of the new isolates showed that cells usually had typical ancyromonad traits, e.g., a laterally compressed, bean-shaped cell with an anterior rostrum and two heterodynamic flagella. With some exceptions, cells from all strains were ~4 ± 0.5 µm long, and ~1.4-1.5 times as long as wide. Most strains broadly resembled *Ancyromonas sigmoides* B-70 (regarded as representative of the first-described species of ancyromonads: Heiss et al., 2010) in that they wobbled rapidly, with varying degrees of amplitude (i.e., cells’ rotation was over sometimes wide and sometimes small angles) and were anchored to the substrate by the distal ends of their posterior flagella, often giving the impression of something akin to gliding motility. This movement is different from gliding as generally understood; in the interests of accuracy and clarity, we hereafter refer to this typical ancyromonad movement as “twitch-yanking”.

All ancyromonad cells that we observed appeared most often to lay on their right side on the substrate (reckoning the posterior flagellum as emerging ventrally). The slender and short anterior flagellum emerged from a subtle depression at the anterior end of the cell. In *Ancyromonas, Caraotamonas* gen. nov., *Striomonas* stat. nov., *Nutomonas, Nyramonas* gen. nov., and *Olneymonas* gen. nov., the anterior flagellum was entirely acronematic, making it uniformly thin and thus much more difficult to see by LM than the anterior flagella in *Planomonas* and *Fabomonas*. The longer posterior flagellum was about twice the length of the cells and held straight posteriorly during linear movement along the substrate.

The representative morphology for all strains is shown in Figures 1–10, and morphological measurements of the cells are summarised in Table 2. Videos showing the movement pattern of all the strains can be found in Movies S1-18.

**FIGURE 1.**
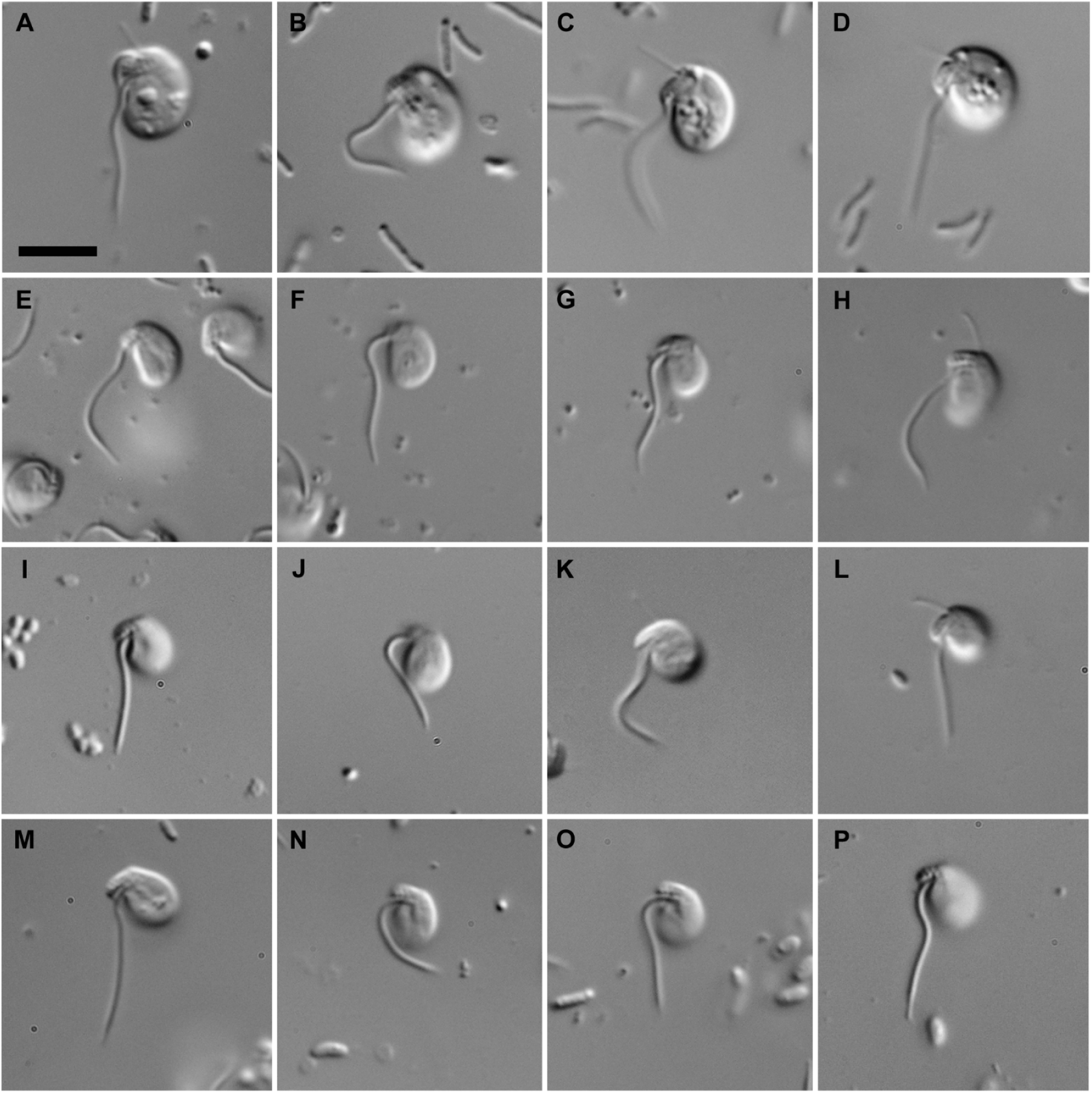
Differential interference contrast light micrographs of living *Ancyromonas* species. **A-D**. *Ancyromonas sigmoides* strain B-70. **E-H.** *Ancyromonas kenti* strain EDM11b. **I-L.** *Ancyromonas kenti* strain AKRosko2018. **M-P.** *Ancyromonas* sp. strain JF-An-6. Anterior of all cells is toward top of page, ventral toward the left. Scale bar = 5 µm.

**FIGURE 2.**
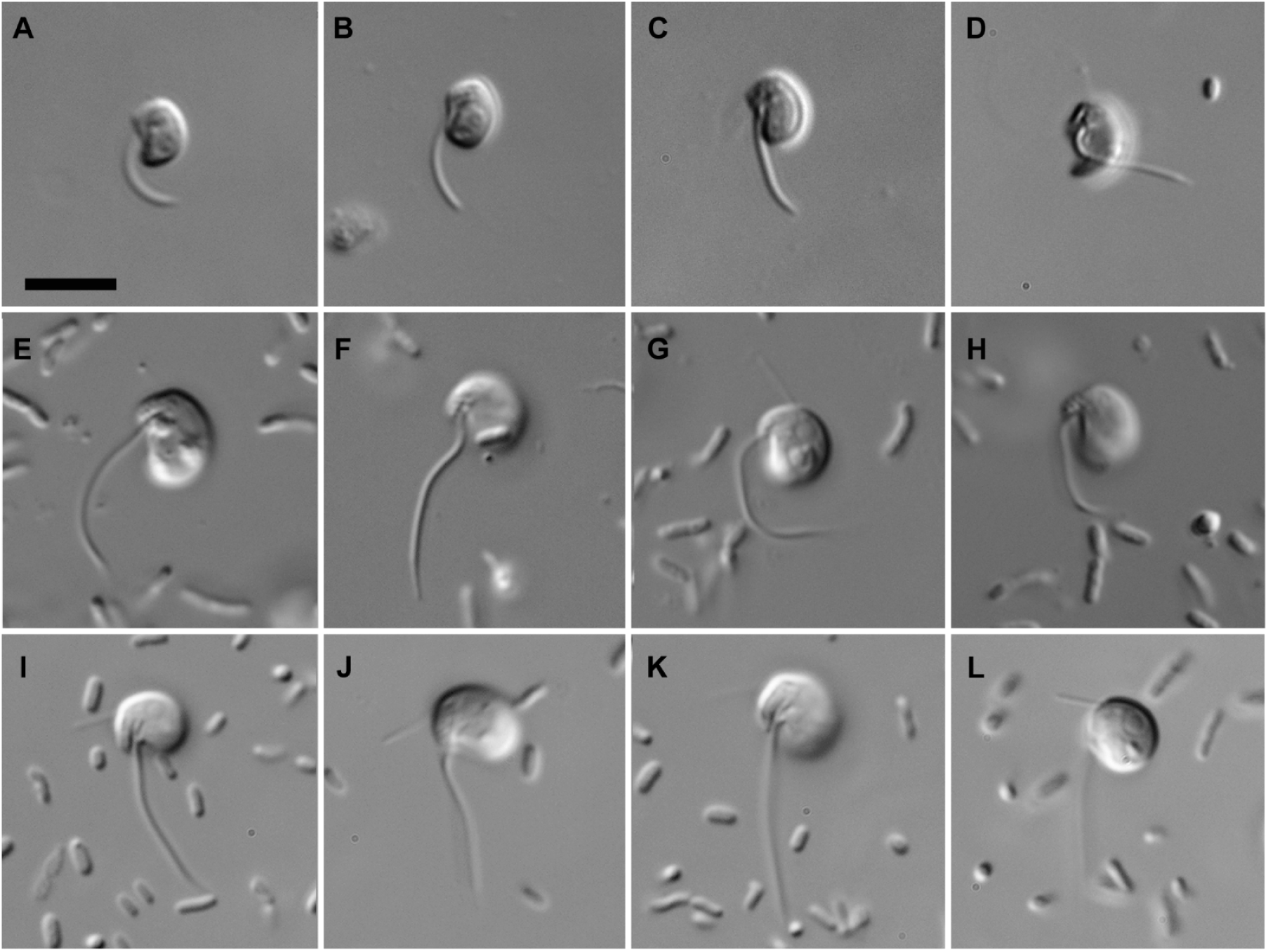
Differential interference contrast light micrographs of living *Ancyromonas* and *Caraotamonas* species. **A-D**. *Ancyromonas mediterranea* sp. nov., strain C362. **E-H.** *Ancyromonas pacifica* sp. nov., strain SRT303. **I-L**. *Caraotamonas croatica* gen. et sp. nov., strain Cro19S5. Anterior of all cells is toward top of page, ventral toward the left. Scale bar = 5 µm.

**FIGURE 3.**
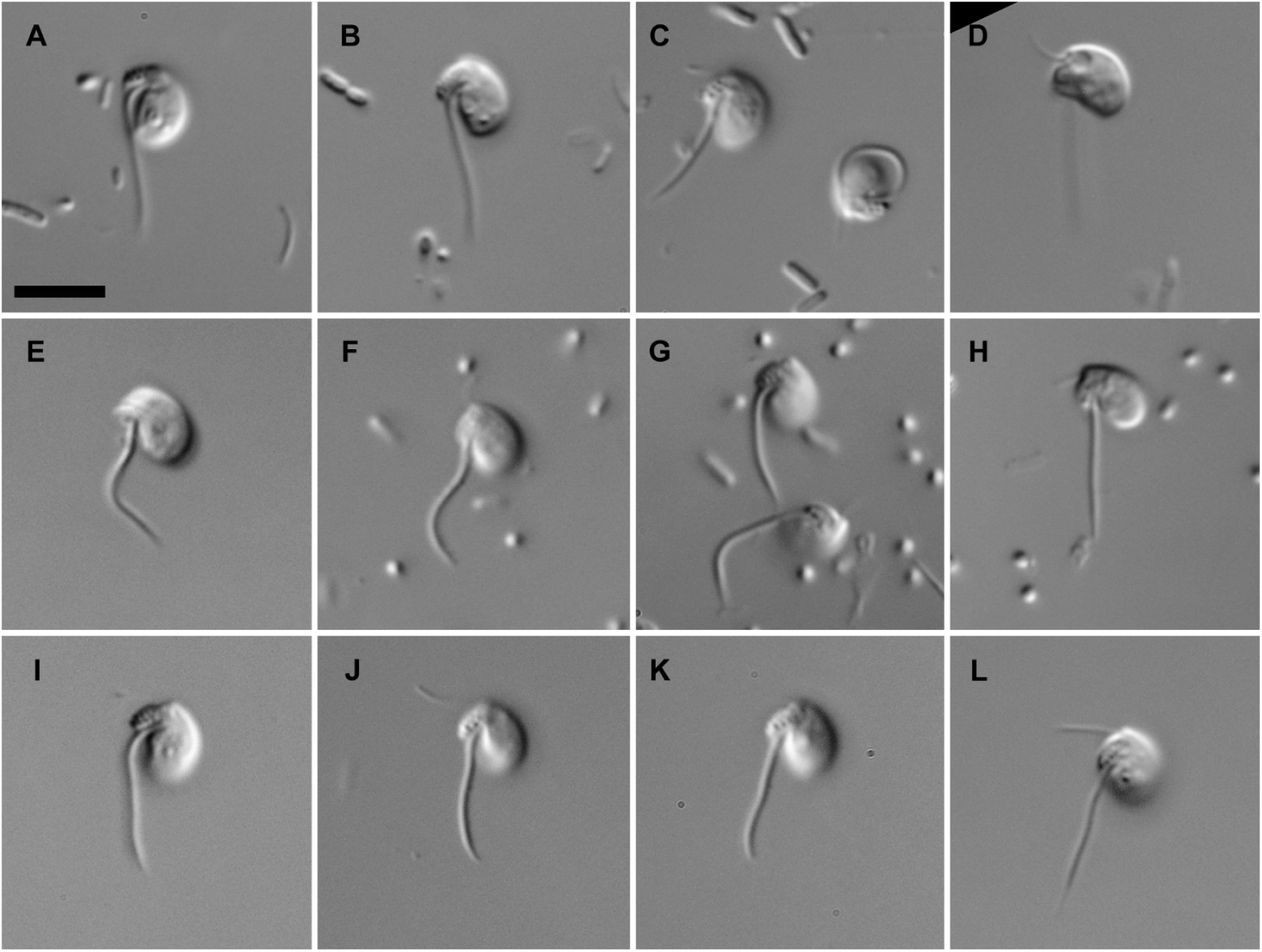
Differential interference contrast light micrographs of living *Nutomonas* and *Nyramonas* species. **A-D**. *Nutomonas limna terrestris* strain OrsayFeb19Ancy. **E-H**. *Nyramonas glaciarum* gen. et sp. nov., strain OrsLand19S1. **I-L**. *Nyramonas silfraensis* gen. et sp. nov., strain OrsLand19S2. Anterior of all cells is toward top of page, ventral toward the left. Scale bar = 5 µm.

**FIGURE 4.**
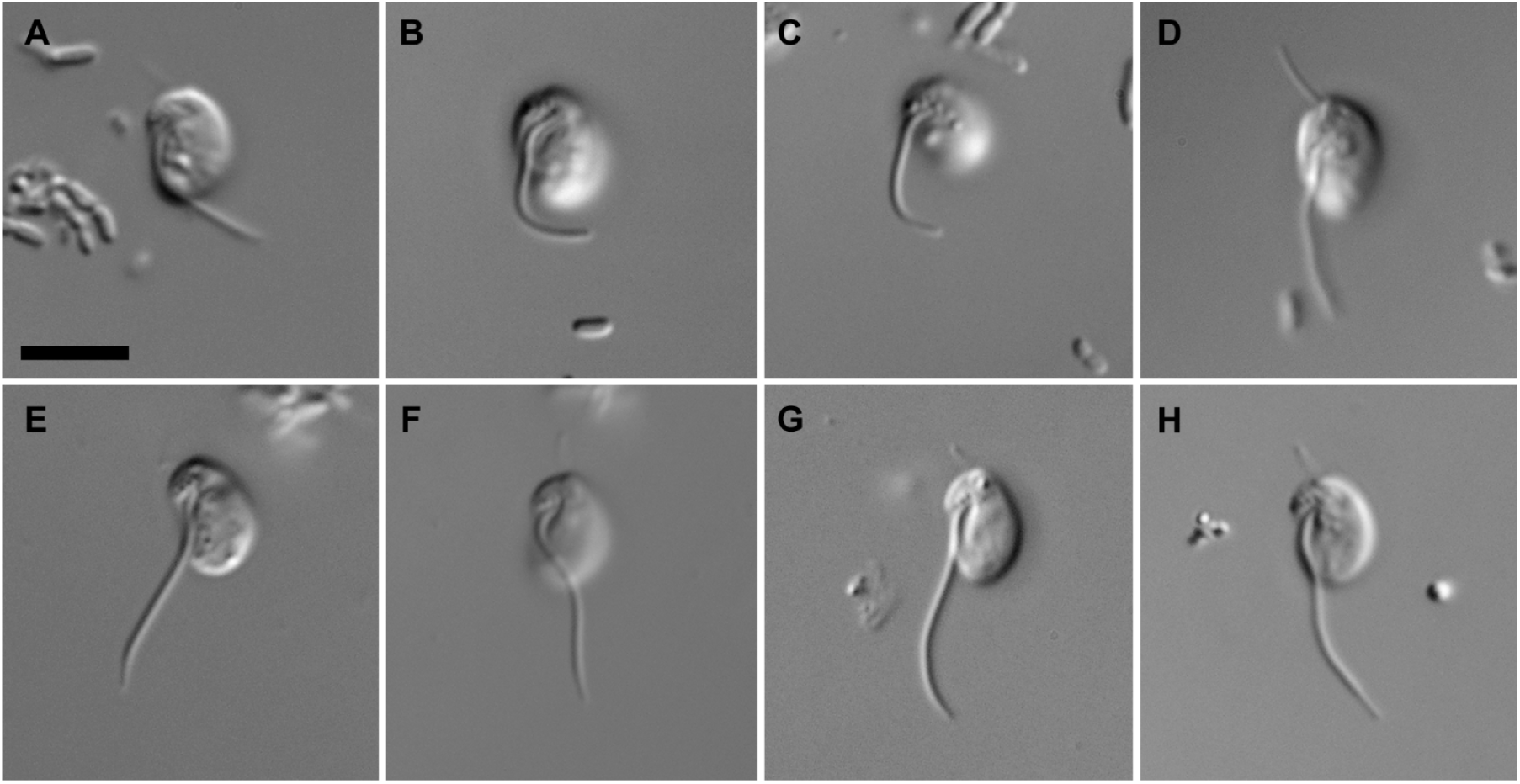
Differential interference contrast light micrographs of living *Olneymonas* and *Striomonas* species. **A-D**. *Olneymonas thurstoni* gen. et sp. nov., strain STRT. **E-H** *Striomonas longa* stat. nov., formerly *Nutomonas*, strain ncfw. Anterior of all cells is toward top of page, ventral toward the left. Scale bar = 5 µm.

**FIGURE 5.**
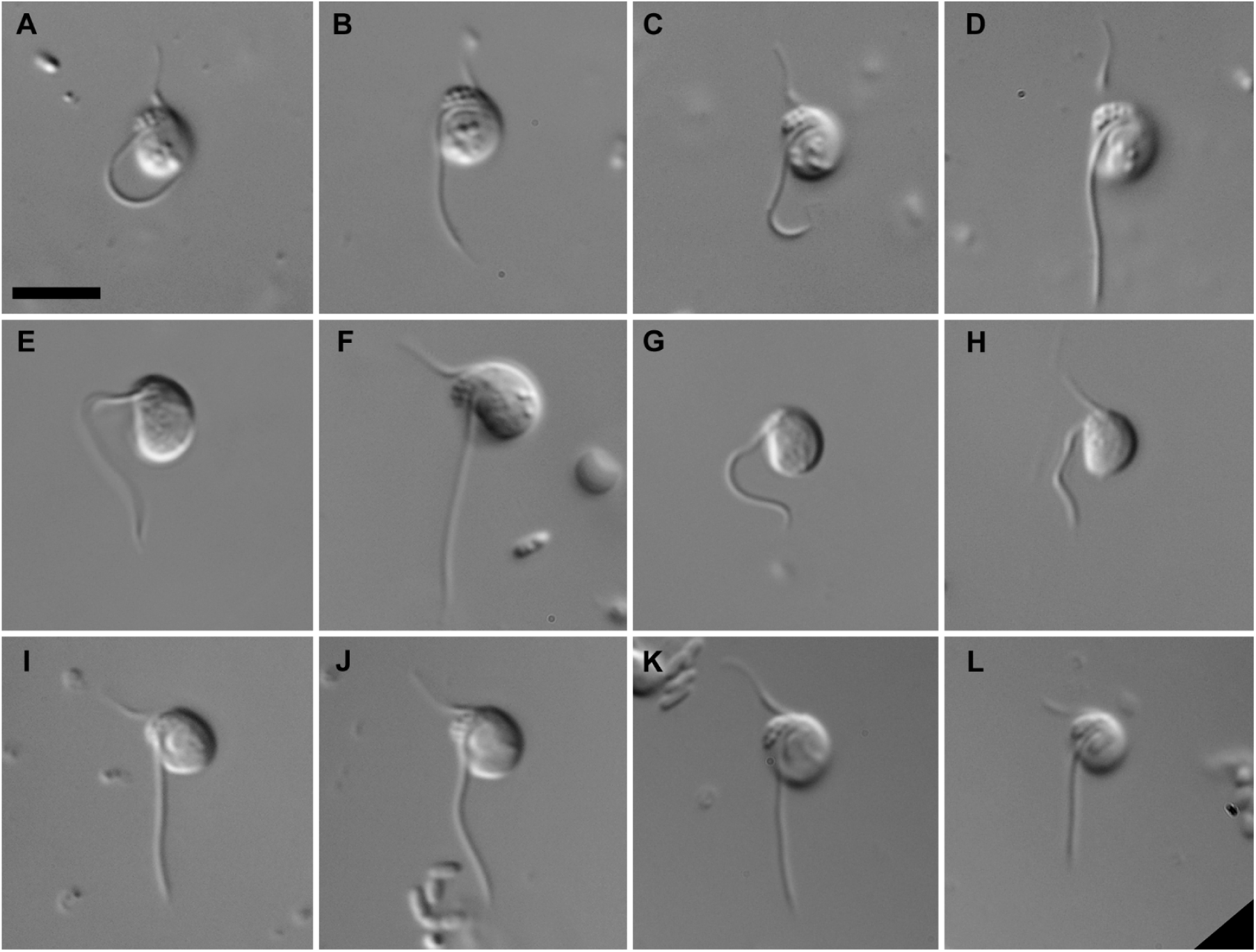
Differential interference contrast light micrographs of living *Planomonas* species. **A-D**. *Planomonas* sp. strain EK-PEI. **E-H**. *Planomonas micra* strain PMRosko2018. **I-L**. *Planomonas* sp. strain SRT307. Anterior of all cells is towar d top of page, ventral toward the left. Scale bar = 5 µm.

**FIGURE 6.**
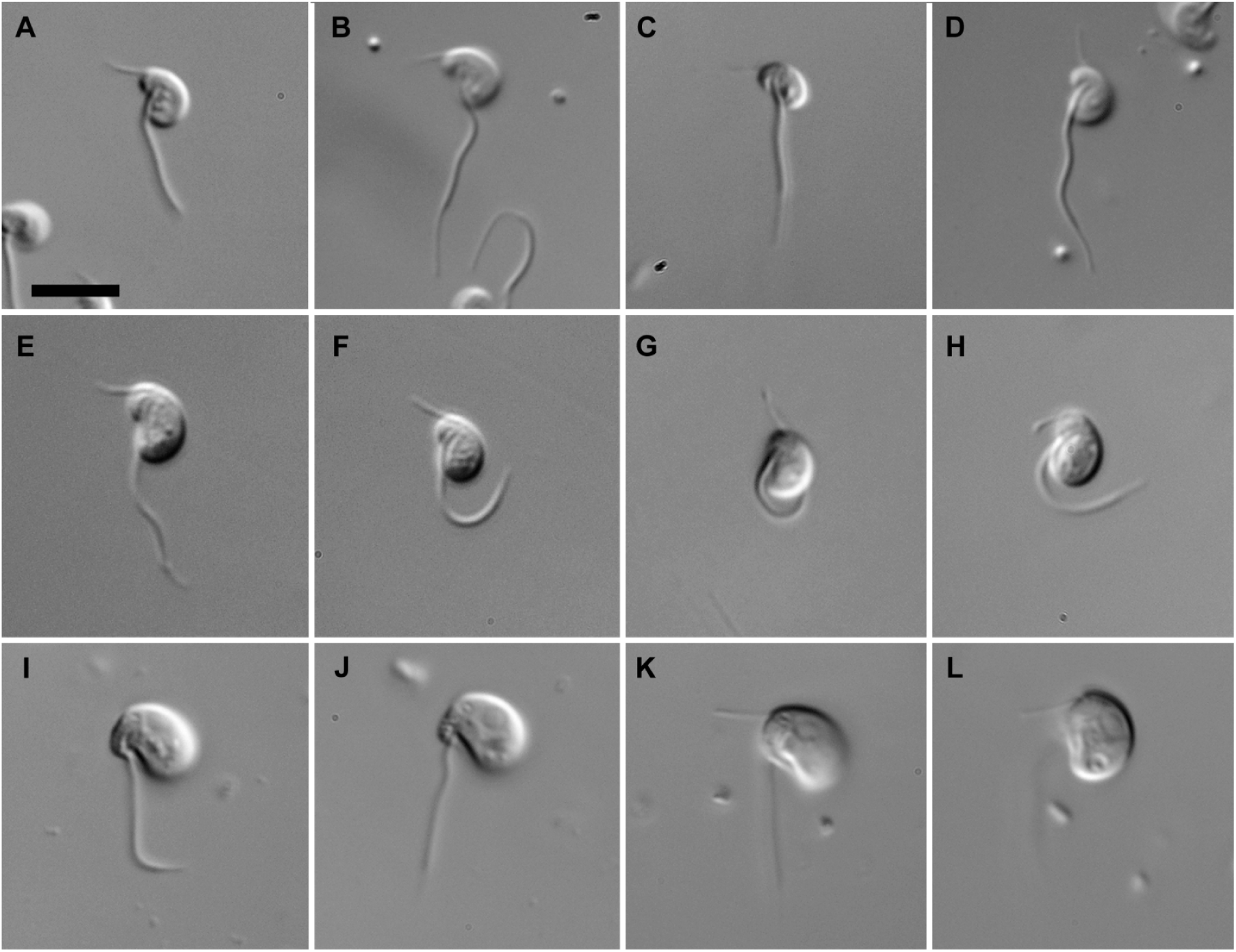
Differential interference contrast light micrographs of living *Fabomonas* species. **A-D**. *Fabomonas tropica* strain NYK3C. **E-H**. *Fabomonas mesopelagica* sp. nov., strain A153. **I-L**. *Fabomonas* sp. strain SRT009. Anterior of all cells is toward top of page, ventral toward the left. Scale bar = 5 µm.

**FIGURE 7.**
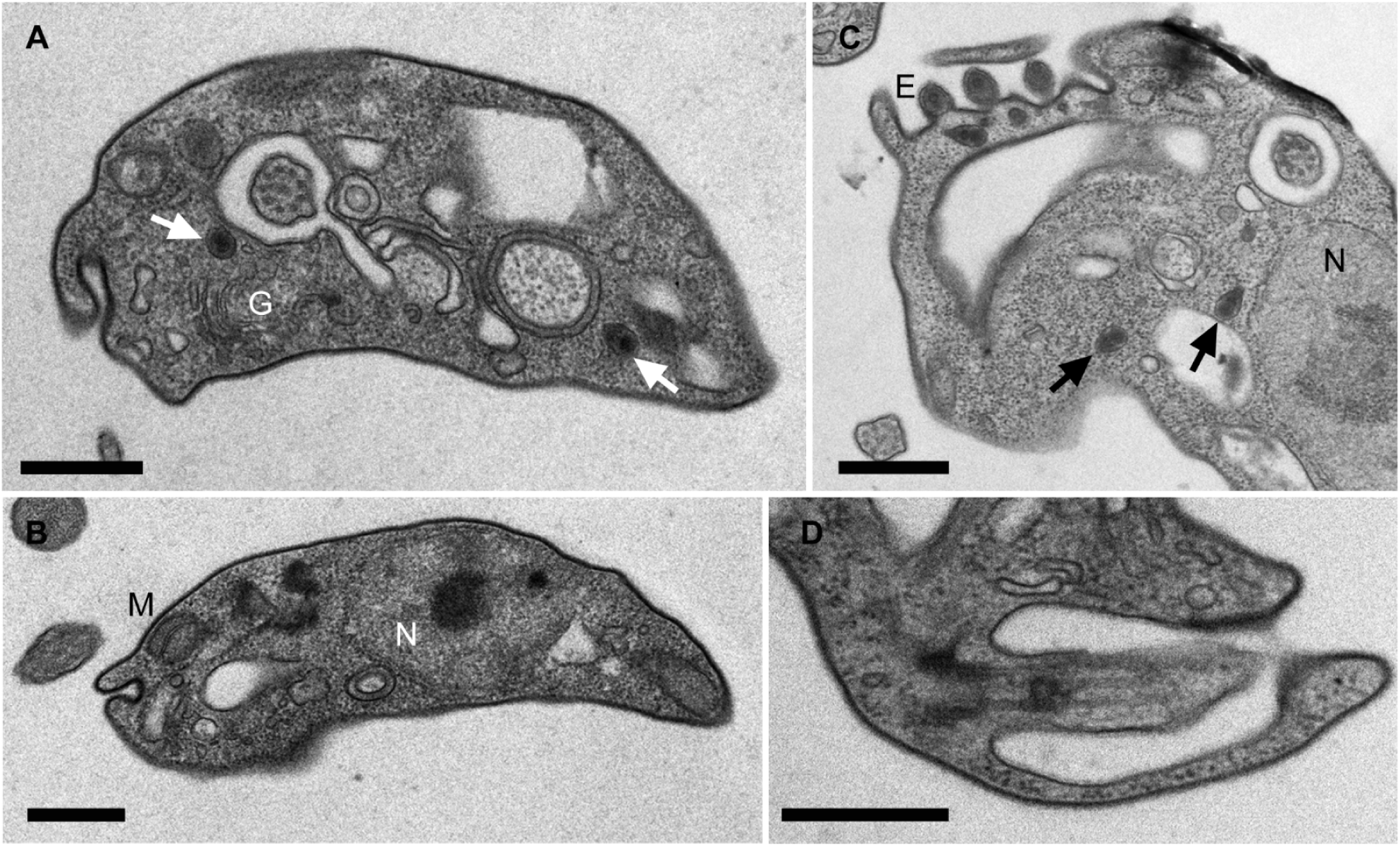
Transmission electron micrographs (TEM) showing general morphology of *Caraotamonas croatica* gen. et sp. nov., strain Cro19S5. **A.** Longitudinal section of cell, with cross section through posterior flagellum, Golgi body (G) and developing extrusomes (arrows). Densely-staining pellicle material is visible just beneath cell surface. **B.** Longitudinal section showing nucleus (N) with conspicuous nucleolus and mitochondrion (M). **C.** Oblique section showing nucleus (N), cross section through anterior flagellum, and row of extrusomes. Arrows indicate developing extrusomes. **D.** Longitudinal section of anterior flagellum. Scale bars = 500 nm in **A-C**, 400 nm in **D**.

**FIGURE 8.**
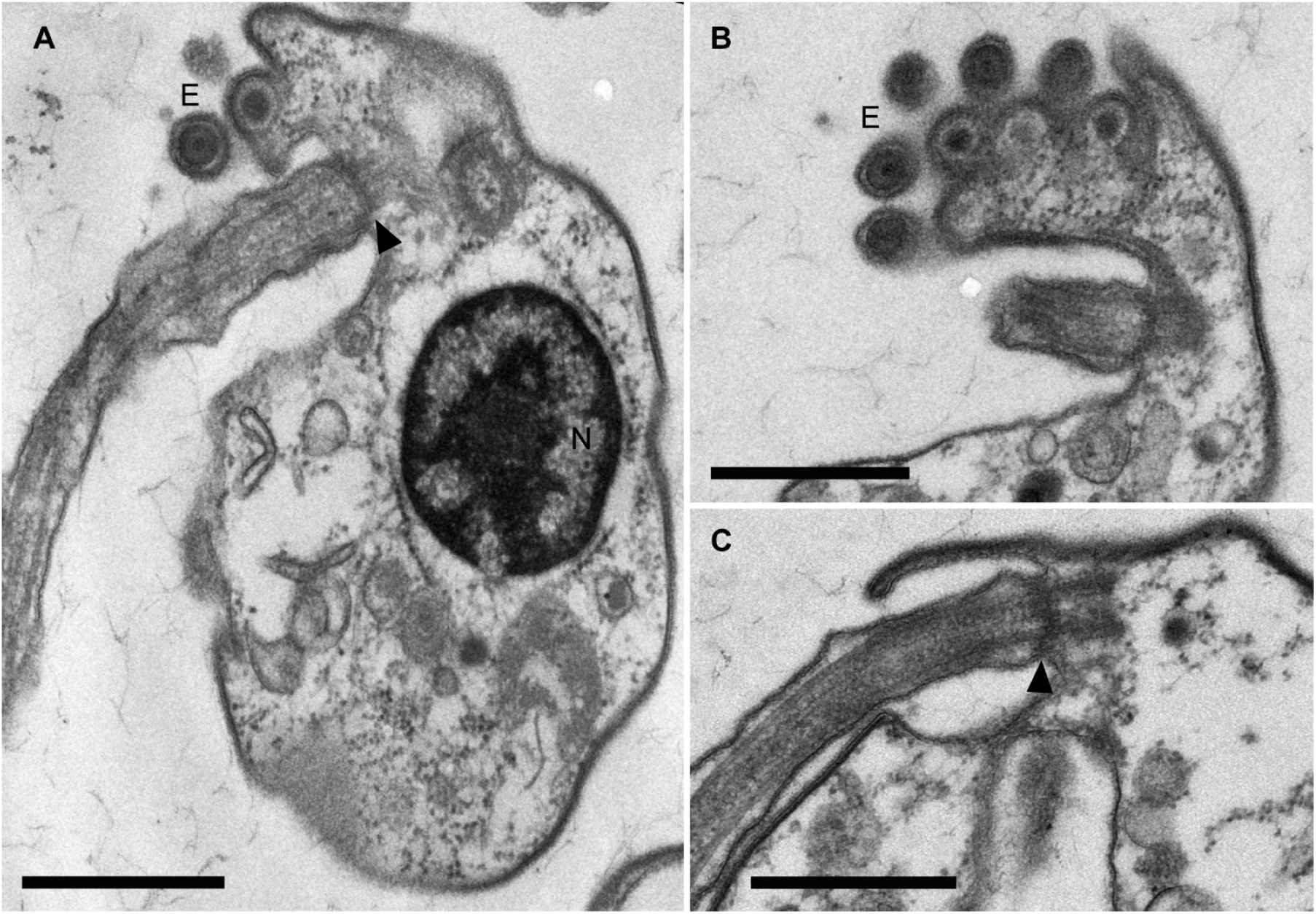
Transmission electron micrographs (TEM) of longitudinal sections showing the general morphology of *Fabomonas mesopelagica* sp. nov., strain A153. **A.** Section showing the nucleus (N), extrusomes (E) and the posterior flagellum. Arrowhead shows the flagellar transition zone. **B.** Section through three rows of extrusomes (E). **C.** Section showing the flagellar transition zone (arrowhead). Scale bars = 500 nm.

**FIGURE 9.**
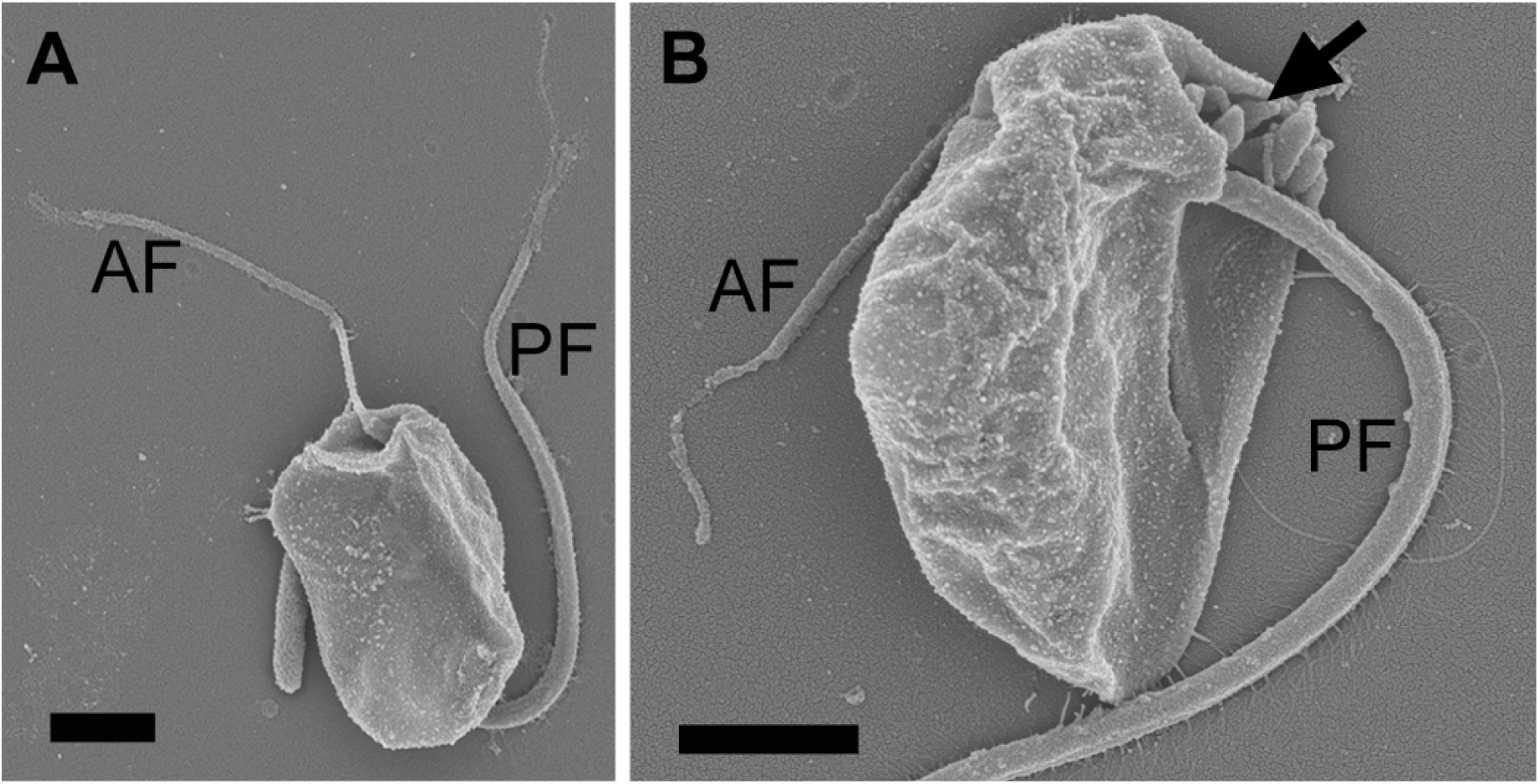
Scanning electron micrographs (SEM) of *Nyramonas silfraensis* gen. et sp. nov., strain OrsLand19S2. **A.** Dorsal side of *Ny. silfraensis* showing a long but strongly acronematic anterior flagellum (AF) and posterior flagellum (PF). **B.** Ventral side of *Ny. silfraensis* showing extrusomes (arrow). Note very fine mastigonemes on PF, projecting away from cell. The anterior of the cells is toward top of page in both images. Scale bar = 1 µm.

**FIGURE 10.**
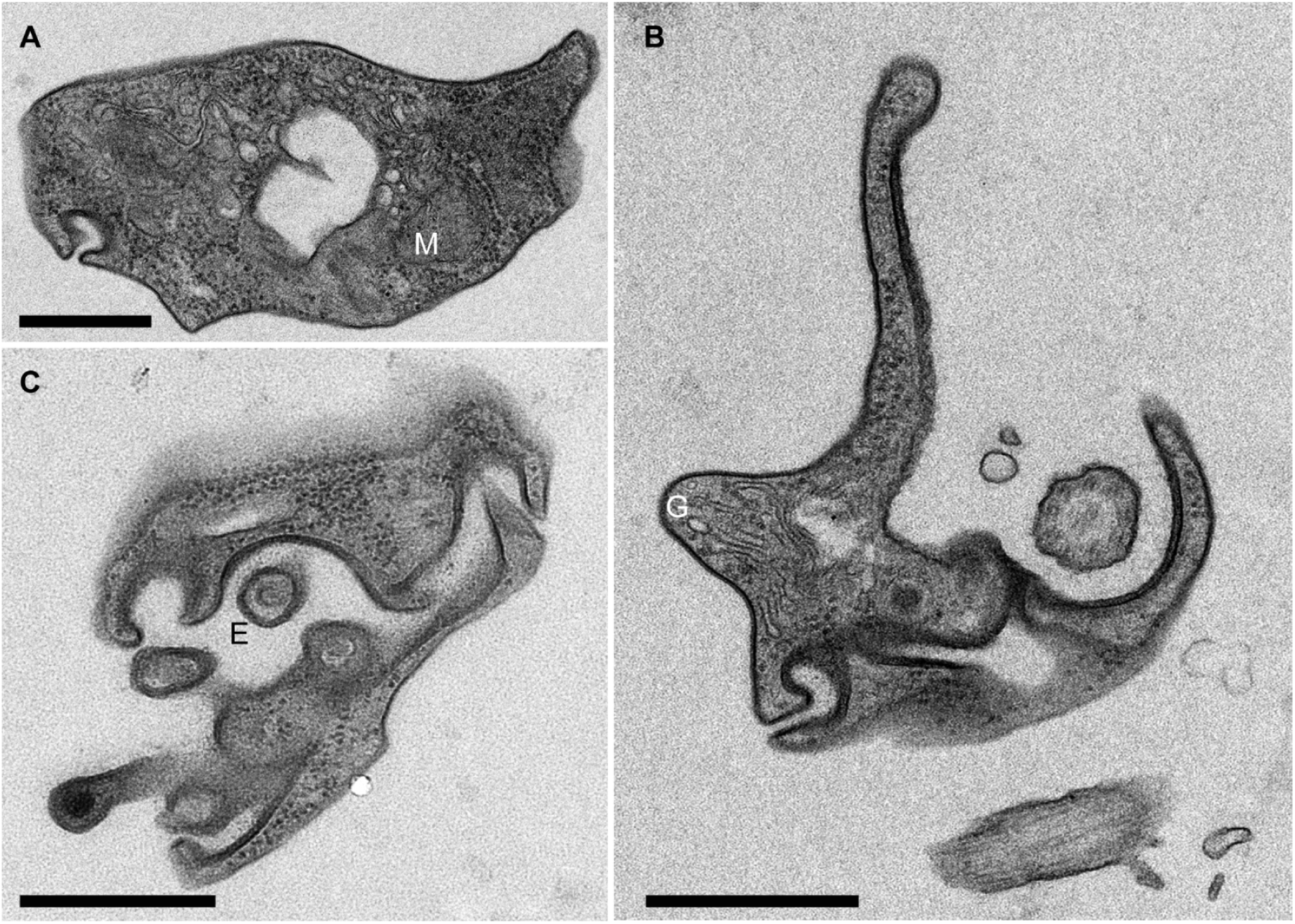
Transmission electron micrographs (TEM) showing the general morphology of *Nyramonas silfraensis* gen. et sp. nov., strain OrsLand19S2. **A.** Section of the posterior part of the cell, showing a mitochondrion (M). Densely-staining pellicle material is visible just beneath the cell the surface. **B.** Section of the anterior part of the cell, showing the Golgi body (G). **C.** Section of the anterior part of the cell, showing putative extrusomes (E). Scale bars = 500 nm.

**Table 2.** Biometric characteristics of the studied strains by light microscopy. Average ± standard deviation given for cell length, width, length/width ratio and length of posterior flagellum.

| Taxonomic name | Isolate | Cell length ($\mu\text{m}$) | Range of cell length ($\mu\text{m}$) | Cell width ($\mu\text{m}$) | Range of cell width ($\mu\text{m}$) | Length/width ratio | Range of Length/width ratio | Posterior flagellum ($\mu\text{m}$) | Range of Posterior flagellum ($\mu\text{m}$) |
| --- | --- | --- | --- | --- | --- | --- | --- | --- | --- |
| *Ancyromonas sigmoides* | B-70 * | 5.4 ± 0.5 | 4.4 - 6.2 | 3.8 ± 0.4 | 2.9 - 4.7 | 1.4 ± 0.1 | 1.3 - 1.8 | 9.2 ± 1.1 | 7.2 - 11.8 |
| *Ancyromonas kenti* | EDM11b * | 4.2 ± 0.5 | 3.2 - 5.1 | 2.9 ± 0.3 | 2.1 - 3.8 | 1.4 ± 0.1 | 1.2 - 1.8 | 7.4 ± 1.2 | 4.1 - 9.4 |
| *Ancyromonas* sp. | JF-An-6 | 4.3 ± 0.5 | 3.3 - 5.4 | 2.9 ± 0.3 | 2.2 - 3.5 | 1.5 ± 0.1 | 1.3 - 1.7 | 7.7 ± 1.1 | 5.1 - 9.2 |
| *Ancyromonas kenti* | AKROSKO2018 | 3.7 ± 0.4 | 3.0 - 4.4 | 2.7 ± 0.3 | 2.2 - 3.2 | 1.4 ± 0.1 | 1.1 - 1.8 | 6.6 ± 0.9 | 5.0 - 8.6 |
| *Ancyromonas mediterranea* sp. nov. | C362 | 3.9 ± 0.4 | 3.3 - 4.5 | 2.3 ± 0.3 | 1.7 - 2.8 | 1.7 ± 0.2 | 1.5 - 1.9 | 7.0 ± 0.7 | 5.8 - 9.1 |
| *Ancyromonas pacifica* sp. nov. | SRT-303 | 4.5 ± 0.5 | 3.6 - 5.5 | 3.0 ± 0.3 | 2.3 - 3.7 | 1.5 ± 0.2 | 1.3 - 2.1 | 8.8 ± 1.4 | 6.6 - 12.0 |
| *Caraotamonas croatica* gen. et sp. nov. | CRO19S5 | 4.5 ± 0.5 | 3.9 - 5.4 | 3.3 ± 0.4 | 2.4 - 4.0 | 1.4 ± 0.1 | 1.2 - 1.7 | 10.4 ± 1.4 | 7.8 - 12.9 |
| *Nutomonas limna* | ORSAYFEB19ANCY | 4.2 ± 0.3 | 3.7 - 4.7 | 3.0 ± 0.3 | 2.3 - 3.4 | 1.4 ± 0.2 | 1.1 - 1.9 | 8.2 ± 1.0 | 6.8 - 10.7 |
| *Striomonas longa* comb. nov. | ncfw * | 4.2 ± 0.9 | 3.3 - 4.9 | 2.7 ± 0.6 | 2.1 - 4.0 | 1.6 ± 0.3 | 1.1 - 1.9 | 8.3 ± 1.9 | 5.5 - 10.4 |
| *Nyramonas glaciarium* gen. et sp. nov. | ORSLAND19S1 | 4.0 ± 0.3 | 3.5 - 4.5 | 2.5 ± 0.2 | 2.0 - 2.9 | 1.6 ± 0.1 | 1.4 - 1.8 | 6.8 ± 0.8 | 5.3 - 8.1 |
| *Nyramonas silfraensis* sp. nov. | ORSLAND19S2 | 4.2 ± 0.8 | 3.5 - 4.9 | 2.9 ± 0.6 | 2.2 - 3.6 | 1.5 ± 0.3 | 1.3 - 1.7 | 7.5 ± 1.5 | 6.0 - 8.9 |
| *Olneymonas thurstoni* gen. et sp. nov. | STRT | 4.9 ± 1.0 | 3.6 - 5.8 | 3.0 ± 0.6 | 2.3 - 3.7 | 1.6 ± 0.3 | 1.3 - 2.1 | 7.3 ± 1.5 | 5.6 - 11.5 |
| *Planomonas micra* | PMROSKO2018 | 3.8 ± 0.4 | 3.1 - 4.7 | 2.8 ± 0.4 | 2.1 - 3.6 | 1.4 ± 0.1 | 1.2 - 1.6 | 7.7 ± 1.8 | 4.7 - 11.4 |
| *Planomonas* sp. | EK-PEI | 4.1 ± 0.4 | 3.4 - 4.8 | 2.9 ± 0.3 | 2.2 - 3.6 | 1.4 ± 0.2 | 1.1 - 1.8 | 8.2 ± 1.3 | 4.9 - 10.4 |
| *Planomonas* sp. | SRT-307 | 3.6 ± 0.4 | 2.9 - 4.4 | 2.6 ± 0.4 | 2.0 - 3.4 | 1.4 ± 0.1 | 1.1 - 1.7 | 8.7 ± 1.3 | 6.2 - 11.5 |
| *Fabomonas tropica* | NYK3C * | 3.5 ± 0.9 | 2.7 - 3.9 | 2.0 ± 0.6 | 1.4 - 2.6 | 1.7 ± 0.5 | 1.4 - 2.4 | 10.6 ± 3.1 | 6.9 - 14.5 |
| *Fabomonas* sp. | SRT-009 | 4.9 ± 1.0 | 4.1 - 5.6 | 3.2 ± 0.6 | 2.7 - 3.5 | 1.6 ± 0.3 | 1.2 - 1.9 | 9.2 ± 1.9 | 7.0 - 10.9 |
| *Fabomonas mesopelagica* sp. nov. | A153 | 3.8 ± 0.4 | 2.9 - 4.4 | 2.6 ± 0.4 | 2.0 - 3.2 | 1.5 ± 0.1 | 1.3 - 1.8 | 9.0 ± 1.6 | 6.0 - 12.9 |
| Previously characterized strains are marked with an asterisk (*). Morphological data of B-70 is taken from Cavalier-Smith et al. (2008) and EDM11b, ncfw and NYK3C are taken from Glücksman et al. (2013). |  |  |  |  |  |  |  |  |  |

Based on a combination of morphological characteristics and molecular phylogeny of the new strains (see below), we identified six strains as belonging to the genus *Ancyromonas,* one strain to the genus *Nutomonas,* three strains to the genus *Planomonas*, and three strains to the genus *Fabomonas*. In addition, we found that seven of our strains did not fit previous descriptions, and accordingly describe three new genera, *Caraotamonas*, *Nyramonas* and *Olneymonas*, and seven new species of ancyromonads belonging to both newly and previously described genera: *A. mediterranea*, *A. pacifica*, *C. croatica*, *Ny. glaciariu*m, *Ny. silfraensis, O. thurstoni,* and *F. mesopelagica* spp. nov. We also transferred *Nu. longa* to the genus *Striomonas* stat. nov. (see Taxonomic Summary).

#### Morphology of *Ancyromonas* isolates

The genus *Ancyromonas* exhibited a wide range in size, including both the largest and some of the smallest cells amongst all of the ancyromonad genera. The anterior end of the cell could be distinguished from the dorsal surface by a curved right-to-obtuse angle between the two surfaces (although the curve was broad enough to mask this in some individual cells of all strains). Cells’ posterior flagella were usually acronematic at the distal end; their anterior flagella were entirely acronematic or nonemergent, and very difficult to see, even under high-power LM. Anterior flagella were practically invisible when live cells were observed. Below, we describe six strains, including four new isolates. These represent two new species, as well as a new strain of the previously described species of *Ancyromonas kenti* (AKRosko2018), and another strain that we did not identify beyond genus (JF-An-6). *Ancyromonas sigmoides*, strain B-70, is the neotype strain of the genus

*Ancyromonas* (Heiss et al., 2010), and originated from the White Sea. Our morphological observations were congruent with the ones from Heiss et al. (2010; 2011), although Cavalier-Smith et al. (2008) reported larger cells for the same strain. Nevertheless, *A. sigmoides* cells were the largest of the ancyromonads that we observed. Uniquely amongst ancyromonads, the anterior of the cell was almost invariably slightly but notably concave (consistent with previous observations; Figure 1A-D; Movie S1).

*Ancyromonas kenti*, strains AKRosko2018 and EDM11b. Overall, members of this species were of sizes and proportions typical of the ancyromonads that we surveyed. All strains’ anterior ends were flattened. Strain EDM11b was isolated from the Irish Sea coast of England, and is the type strain of this species (Glücksman et al., 2013). Our observations were congruent with the previous morphological characterisation of this strain (Figure 1E–H; Movie S2). Strain AKRosko2018 was isolated from an intertidal sediment sample in Roscoff, France. Although the size range of this strain’s cells overlapped with those of other *Ancyromonas* species, it was somewhat smaller than its conspecifics. In addition, the posterior flagellum’s acroneme tended to be shorter and less obvious than in other strains (Figure 1M–P; Movie S4).

*Ancyromonas* sp., strain JF-An-6. Although similar to *A. kenti* morphologically, and branching with it in our 18S rRNA gene trees (see below), lack of ITS sequence data prevented us from confirming this strain’s identity as *A. kenti*. It was isolated from Bradley Beach, Neptune Township, New Jersey, USA. Its posterior flagella were almost always visibly acronematic (Figure 1I–L).

*Ancyromonas mediterranea* sp. nov., strain C362. This strain was isolated from the mesopelagic water column (250 m deep) in Villefranche Bay, France. Cells were notably elongated compared to most other ancyromonads, and in general smaller than other ancyromonads as a whole, although the range of their size overlapped. The anterior end of the cell was flattened, as with *A. kenti*, and the posterior flagellum was usually visibly acronematic (Figure 2A–D; Movie S5).

*Ancyromonas pacifica* sp. nov., strain SRT303. The characteristic trait of this new species was its slow movement. As with other ancyromonads, cells anchored themselves to the substrate by the posterior flagellum, but exhibited a slow swing from side to side rather than the flicking movement characteristic of other ancyromonads. They moved forward very slowly while attached to the substrate. However, members of this strain had the same rapid “twitch-yanking” movement as other ancyromonads when cultured in different media (specifically, Erd-Schreiber medium supplemented with 3% Cerophyl). As with *A. kenti* and *A. mediterranea*, cells’ anterior ends were flattened. Digestive vacuoles were observed in the middle or rear end of the cell and contained bacteria (Figure 2E–H; Movie S6).

#### Morphology of *Caraotamonas croatica* gen. et sp. nov., strain Cro19S5

This strain was isolated from a saltwater lake in Mljet Island, Croatia. The transition between the dorsal and anterior surfaces was more consistently rounded than in any species of *Ancyromonas*, as was also the case for the transition between the ventral and posterior surfaces. Cells’ anterior flagella were entirely acronematic and difficult to observe, as with *Ancyromonas* species. The distal end of the posterior flagellum was also acronematic (Figure 2I–L). When moving on the substrate, cells of *Caraotomonas* did not change direction as much as those of *A. sigmoides*, and the amplitude of their oscillation was consistently small (Movie S7).

#### Morphology of *Nutomonas limna terrestris*, strain OrsayFeb19Ancy

This strain was isolated from a shallow freshwater ditch in the University of Paris-Saclay campus, France. The anterior flagella of these cells were entirely acronematic, and usually difficult to see by LM, as in those of *Ancyromonas* and *Caraotamonas* species (Figure 5B). Cells of this strain matched the description of *Nutomonas limna* made by Cavalier-Smith et al. (2008), including their freshwater habitat. A contractile vacuole was observed near the insertion of the anterior flagellum.

#### Morphology of *Nyramonas* gen. nov

We assigned two freshwater strains, OrsLand19S1 and OrsLand19S2, to the new genus *Nyramonas*. Their size range overlapped with those of *Nutomonas* species, but cells were moderately elongated. In addition, while the anterior surface of the cell was flattened as with other freshwater ancyromonads, the angle between the anterior and dorsal surfaces of the cell was right to acute. The anterior flagellum of both strains was long, typically only slightly shorter than the cell length, but acronematic throughout its visible length, and thereby difficult to see by light microscopy. The contractile vacuole was situated near the base of the anterior flagellum. Both strains were isolated from Iceland: OrsLand19S1 (*Ny. glaciarum* sp. nov.: Figure 3E–H; Movie S10) from the Sólheimajökull glacier, and OrsLand19S2 (*Ny. silfraensis* sp. nov.: Figure 3I–L; Movie S11) from the Silfra rift.

#### Morphology of *Olneymonas thurstoni* gen. et sp. nov., strain STRT

This strain was isolated from the surface of a rainbow trout (*Oncorhynchus mykiss*) caught in Olney Pond, Lincoln, RI, USA. *Olneymonas thurstoni* cells were distinctly larger than those of *Nyramonas* and *Nutomonas* species, and the anterior and dorsal surfaces of the cell were distinguished by an obtuse angle. As with *Caraotamonas*, cells’ oscillation was over a consistently narrow angle. An acronematic distal end to the posterior flagellum was often observed. A contractile vacuole was observed near the insertion of the anterior flagellum. The posterior flagellum length was highly variable in length, from 5.6 to 11.45 µm long (7.3 ± 1.5 µm) (Figure 4A–D; Movie S12).

#### Morphology of *Striomonas longa* stat. nov., strain ncfw (CCAP 1958/5)

This strain was originally described as *Nutomonas (Striomonas) longa* by Glücksman et al. (2013). However, the strain’s 18S rRNA gene sequence was quite divergent from that of its congeners (Figure 11), and it was additionally distinguished by its elongated morphology. We also observed this distinctive morphology, and noted as well that cells had a right-to-acute angle between the anterior and dorsal surfaces of the cell (unlike other *Nutomonas* spp.), although the cells in our strain measured slightly smaller and were much more rounded than those in the original description (3.5-5.0 µm long, 3.0-3.5 µm wide, ~1.31 ratio: Glücksman et al., 2013) (Figure 4E–H; Movie S9). Given these differences in morphology, and that the phylogenetic placement of this species as sister to other genera rendered the genus *Nutomonas* paraphyletic (Figure 11), we propose to raise *Striomonas* to the rank of genus.

**FIGURE 11.**
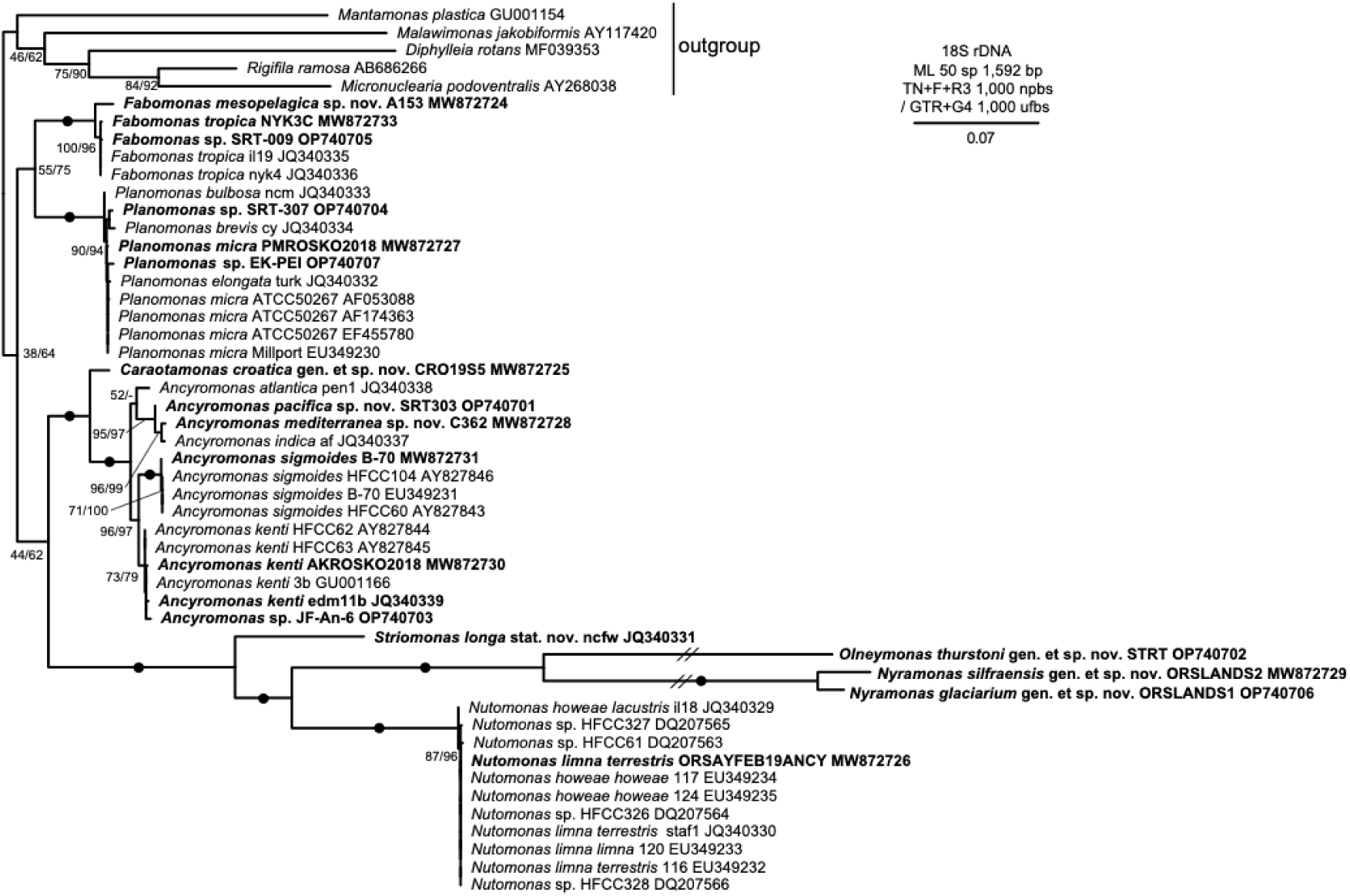
18S rDNA phylogenetic tree of 45 characterised ancyromonads and five outgroups. Maximum likelihood tree inferred from 1,592 nucleotides using TN+F+R3 (the best fitting model as per ModelFinder implemented in IQ-TREE). Support values correspond to percentages of 1,000 replicates of nonparametric bootstraps under TN+F+R3 model and ultrafast bootstraps under GTR+G4 model, respectively. Strains analysed in this study are in bold. Filled circles show maximally supported clades. Long branches (//) reduced by factor of 2. Scale bar indicates the number of expected substitutions per site.

#### Morphology of *Planomonas* isolates

We isolated three strains that we identified as members of the genus *Planomonas*. As per Glücksman et al. (2013), *Planomonas* and *Fabomonas* differ from other ancyromonads in having an anterior flagellum that is not entirely acronematic (Figs. 5A, C, K), and often readily visible under the light microscope. Cells of *Planomonas* were generally rounder than those of *Fabomonas*; this included the transition between the anterior and dorsal surfaces of the cells, which tended to be more angular in strains identified as *Planomonas* species.

Strain EK-PEI was isolated from Prince Edward Island, Canada. (Figure 5A–D; Movie S13). PMRosko2018 (Figure 5E–H; Movie S14) was isolated from Roscoff, France, and strain SRT307 (Figure 5I–L; Movie S15) was isolated from the skin of a barracuda caught in a lagoon in Iriomote Island, Japan. On average, cells of PMRosko2018 were slightly larger than those of SRT307 and smaller than those of EK-PEI. In addition, SRT307’s posterior flagella tended to be slightly longer than those of each of our other two strains of *Planomonas*, and their anterior flagella appeared to have longer non-acronematic regions than those of EK-PEI (cf Figure 5A, C, K).

#### Morphology of *Fabomonas* isolates

We observed three strains, including the type strain of the genus, *Fabomonas tropica* NYK3C.

*Fabomonas tropica*, strain NYK3C. Our morphological observations of this strain were congruent with the original characterisation (Glücksman et al., 2013). Cells of this strain were the most elongated of all ancyromonad strains that we observed. The posterior flagellum of this strain was also the longest among our observed strains (Figure 6A–D; Movie S16).

*Fabomonas mesopelagica* sp. nov., strain A153. This strain was isolated from 250 m below the surface of the Mediterranean Sea, off the coast of Villefranche, France. Cells were smaller and rounder than those of *F. tropica* (Figure 6E–H; Movie S17).

*Fabomonas* sp., strain SRT009. The motility of the SRT009 strain was different from that of any other ancyromonad species, and due to the lack of ITS data, it was not assigned up to a species level. The cell usually lay on the substrate with the posterior flagellum swinging alongside the cell body as they moved forward together. As with the unusual movement observed for *Ancyromonas pacifica*, though, this could be a result of the media in which the cells were cultured; when cultured in a mixture of freshwater Cerophyl and natural seawater (specifically, 10% freshwater Cerophyl, 50% natural seawater, 40% distilled water), members of this strain exhibited normal ancyromonad “twitch-yanking” movement. In addition, cells of this strain were markedly larger than those of NYK3C, and differed as well from that strain in being of typical ancyromonad proportions. They also tended to have somewhat concave anterior ends. Their anterior flagella were shorter than the cell length, and hard to observe under the light microscope (Figure 6I–L; Movie S18).

### Electron microscopy

All specimens from new genera were observed under transmission electron microscopy (TEM), and presented a single layer of densely-staining pellicle material appressed to the plasma membrane. This was seen beneath the entire cell surface except for pockets at the bases of both anterior and posterior flagella (e.g., Figure 7A). We did not observe any ‘stacked membrane’ structures similar to those reported in *Ancyromoans sigmoides* (B-70) (Heiss et al., 2011) in any of our preparations.

#### Ultrastructure of *Caraotamonas croatica* gen. et sp. nov., strain Cro19S5

Cells of *Caraotamonas croatica* possessed a nucleus containing a prominent nucleolus (Figure 7B-C). At least two rows of extrusomes were positioned in the rostrum at the anterior side of the cell. Developing extrusomes were observed near the Golgi body and throughout the cell. Axosomes characterised the transition regions between the basal bodies and their associated flagella (Figure 7D).

#### Ultrastructure of *Fabomonas mesopelagica* sp. nov., strain A153

Our TEM observations of *Fabomonas mesopelagica* suggested that the nucleus in this species was located in the anterior region of the cell, close to the two basal bodies (Figure 8A). At least three rows of extrusomes were positioned in the rostrum (Figure 8B), and the outer edge of the flagellar pocket appeared extremely thin, comprising only the cell membrane and a layer of pellicle along its outside face (Figure 8C). Transitional plates between the flagella and the basal bodies were observed (Figure 8C).

Surface and interior ultrastructure of *Nyramonas silfraensis* gen. et sp. nov., strain OrsLand19S2.

Scanning electron microscopy (SEM) confirmed that each cell of *Nyramonas silfraensis* gen. et sp. nov., strain OrsLand19S2, had two flagella and a shallow ventral groove (Figure 9). The anterior flagellum of *Ny. silfraensis*, which was extremely hard to observe by LM, was clearly seen under SEM to comprise a thicker basal region extending to the point of emergence from the flagellar pocket and an acronematic region extending for the rest of its length (Figure 9A). The posterior flagellum possessed very thin, simple, ~200-nm-long mastigonemes (thin hairs) (Figure 9B). Several extrusomes were observed at the anterior of the cell, in the region proximate to the point of insertion of the posterior flagellum. TEM shows that *Nyramonas*appears to have loosely packed flat mitochondrial cristae (Figure 10A), and its Golgi body was located at the anterior part of the cell (Figure 10B). At least two rows of extrusomes were situated in the rostrum (Figure 10C).

### 18S rRNA gene phylogeny of ancyromonads

We inferred a 50-taxon phylogeny based on the 18S rRNA gene, which included sequences of 1) our 14 new isolates; 2) the previously described strains B-70 and NYK3C, for each of which we re-sequenced the 18S rRNA genes; 3) 29 ancyromonad strains from public depositories (Glücksman et al., 2011; 2013); and 4) five outgroup taxa, chosen based on recent phylogenomic analyses (Brown et al., 2018): *Mantamonas*, *Diphylleia*, *Rigifila*, and *Micronuclearia* (the ‘CRuMs’), and *Malawimonas*. The resulting tree was largely unresolved for the deep splits within the ancyromonads, and surprisingly without support for ancyromonads as a clade (Figure 11).

However, each ancyromonad genus was recovered, and with maximal support. In addition, several groups of genera formed strongly supported clades: in particular, *Ancyromonas* and *Caraotamonas* grouped together with maximal support, as did all freshwater isolates: *Striomonas*, *Nyramonas*, *Olneymonas*, and *Nutomonas*. Two of our novel strains in the *Ancyromonas* clade (AKRosko2018 and JF-An-6) grouped with previously identified *A. kenti* strains with moderate support. Two other strains (C362 and SRT303) grouped robustly with *A. indica*, and sister (albeit without support) to *A. atlantica*. Within the freshwater clade, *S. longa* was sister to all other members, with maximal support. The genus *Nutomonas* sensu stricto (equivalent to the subgenus *Nutomonas* as defined by Cavalier-Smith in Glücksman et al., 2013), consisting only of the species *Nu. howeae* and *Nu. limna*, was represented by 11 almost identical sequences, including our isolate OrsayFeb19Ancy. Differentiation of species of *Nutomonas* has been determined by analysis of ITS2 sequences (Cavalier-Smith et al., 2008), which identified our strain as *Nu. limna terrestris* (In contrast, 18S phylogeny is congruent with ITS2 sequence types for the genera *Ancyromonas* and *Planomonas*: data not shown). The genus *Nutomonas* branched with maximum support as sister to a clade containing the two new genera *Olneymonas* and *Nyramonas*, each forming a long branch on the tree. The nearly identical sequences from the three isolates PMRosko2018, EK-PEI and SRT307 clustered with all members of the genus *Planomonas.* As with *Nutomonas*, species of *Planomonas* are better differentiated based on ITS2 sequence analysis (Glücksman et al., 2013). This enabled us to identify PMRosko2018 as *P. micra*; however, since we lacked ITS2 sequences for our other two *Planomonas* strains, we could not identify them beyond the genus level. Thus, we regarded EK-PEI and SRT307 as “*Planomonas* sp.”. Finally, the *Fabomonas* clade contained a robust split between one of our new strains, *F. mesopelagica* A153, and a clade that contained four strains of *F. tropica*, including both the type strain, NYK3C, and one new isolate, *Fabomonas* sp. SRT009.

### Survey of environmental ancyromonad sequences

We searched for ancyromonad sequences in the PR2, Silva and GenBank nucleotide databases and retrieved 178 environmental 18S rDNA sequences (Table S2). In addition, we retrieved 146 additional ancyromonad sequences from operational taxonomic units (OTUs) identified in five recent protist metabarcoding studies: three from freshwater (David et al., 2020; 2021; Iniesto et al., 2020) and two from marine environments (Logares et al., 2020; Massana et al., 2015). Finally, we obtained nine additional environmental sequences from previously sampled marine and freshwater environments (Torruella et al., 2017) by PCR amplification of 18S rRNA genes using ancyromonad-specific primer pairs (Table S3), cloning and sequencing. This resulted in a dataset with 333 18S rRNA gene sequences of environmental origin.

We used this environmental 18S rDNA dataset to infer phylogenies using multiple evolutionary models (Figures 11, 12 and S1). The same main clades as those from the 50-taxon phylogeny were recovered, although with lower support, and their basal relationships remained unresolved. However, the clade comprising all cultured freshwater ancyromonads (*Nutomonas*, *Nyramonas*, *Olneymonas*, and *Striomonas*), which also included 106 environmental sequences, was highly supported.

**FIGURE 12.**
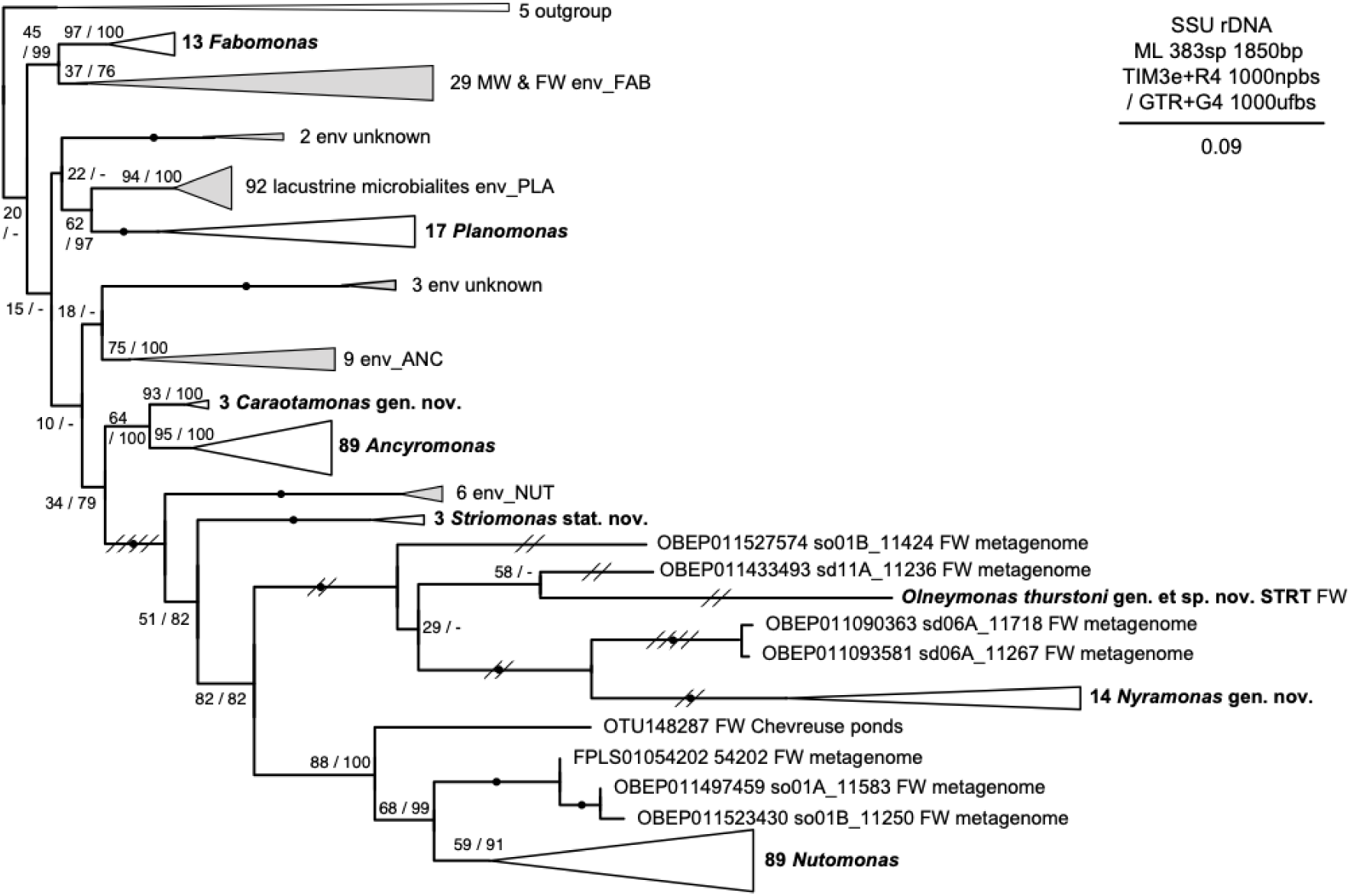
18S rDNA phylogenetic tree of 45 characterised ancyromonads, 333 environmental ancyromonad sequences from public databases and five metabarcoding studies, and five outgroups. Maximum likelihood tree inferred from 1,850 nucleotides using TIM3eR4 model. Support values correspond to percentages of 1,000 replicates of non-parametric bootstraps under TIM3eR4 model and ultrafast bootstraps under GTR+G4 model, respectively. Clades with characterised species are in bold. Colored triangles correspond to highly supported clades of environmental sequences (see Table S2 for details). Long branches reduced by factor of 2 (//) or 4 (////); fully expanded tree shown in Figures S1A and S1B. Scale bar indicates the number of expected substitutions per site.

Interestingly, environmental sequences seemed to represent several novel lineages distinct from the four genera described in this work, and hint at a wider diversity of ancyromonads than is presently known. *Caraotamonas* and *Ancyromonas* branched together with 80 marine environmental sequences, clade env_ANC (Figure S1). These were highly similar, in contrast to the multiple and divergent environmental sequences found in the freshwater clade (e.g., env_NUT).

The env_PLA clade (comprising 92 sequences from freshwater microbialites) branched sister to a clade of *Planomonas* sequences, with moderate support. Finally, and similarly, the env_FAB clade, containing 29 sequences from both freshwater and marine environments, appear as sister to *Fabomonas,* although with moderate statistical support.

## DISCUSSION

### Diversity and ecology of ancyromonads

Ancyromonads have been identified by 18S rRNA gene amplicon-based studies in various ecosystems since early times (Scheckenbach et al., 2005; Stoeck et al. 2006) (Table S2). However, in many public databases they appear classified as “Apusozoa” (Cavalier-Smith 1997). However, “Apusozoa” is not a valid taxon, since it grouped organisms that do not form a monophyletic group (e.g. Brown et al., 2013; Torruella et al., 2023):, ancyromonads plus apusomonads, sometimes even including other lineages, such as breviates, collodictyonids, and mantamonads. Unfortunately, this blurs the identification of bona fide ancyromonads in the numerous eukaryotic metabarcoding studies that have treated the sequences from these clades as a single group (e.g., Singer et al., 2021). Consequently, studies ascribing sequences to “Apusozoa”, without specifying a genus are not informative about the true diversity and distribution of ancyromonads. Such data therefore were not included in our study, and we encourage others to be cautious when using sequence data only identified as “Apusozoa”. Further, we strongly recommend that the name “Apusozoa” be abandoned altogether (as PR2 database version 5 recently announced).

Our phylogenetic trees supported not only the four clades (*Ancyromonas*, *Fabomonas*, *Nutomonas* and *Planomonas*) found by Glücksman et al. (2013) and formally described as different ancyromonad genera, but also several new undescribed genus-level clades, and revealed more cryptic biodiversity than previously known (Figures 11 and 12). In addition, our survey of environmental databases confirmed the cosmopolitan distribution of ancyromonads, which also seem ecologically diverse, being present from deep marine sediments to coastal sites, lakes and brooks. Metabarcoding surveys using general eukaryotic primers seem to be able to uncover parts of ancyromonad biodiversity, but more diversity is revealed when these are complemented by the use of, ancyromonad-specific primers. As an example, *Nutomonas* 18S rDNA sequences were identified from the same three sites using universal eukaryotic primers (David et al., 2021). Using specific primers in the same samples, we recovered the *Nutomonas* sequences and also found a *Nyramonas* sequence (Figure S1). Most likely, this could be due to the lower abundance of the latter in the environment.

Based on our survey of environmental data (Figure 12), we can hypothesize that ancyromonad species were ancestrally marine, with at least three independent transitions to freshwater environments: one at the root of *Striomonas*, *Nyramonas*, *Olneymonas*, and *Nutomonas*, one in a possible sister lineage to *Planomonas*, and another within the env_FAB clade (Figure 12 and S1). Interestingly, the latter contained sequences both from marine and freshwater sources, suggesting that, while most cultured marine ancyromonads are euryhaline (Heiss, pers. observ.), these ancyromonads are able to adapt further, possibly transitioning multiple times between marine and freshwater habitats.

As noted above, none of our trees supported any intergeneric relationships amongst the previously-identified clades. This is in contrast to previous studies (in particular Glücksman et al., 2013), which placed *Fabomonas* as sister either to *Planomonas* or to all other ancyromonads, and which invariably identified *Ancyromonas* and the cultured freshwater clade (then comprising only *Nutomonas* and *Striomonas*) as sisters. Glücksman et al. (2013) formalised these relationships by defining the family Planomonadidae as comprising the genera *Planomonas* and *Fabomonas*, and Ancyromonadidae as containing *Ancyromonas* and *Nutomonas*. If we retain this division, *Caraotamonas*, *Striomonas*, *Nyramonas*, and *Olneymonas* would then also be assigned to the family Ancyromonadidae. However, in light of our findings, it seems that the relationships amongst ancyromonad clades are difficult to resolve based only on 18S rRNA gene phylogenetic analyses. To uncover those deep relationships and reconstruct a robust tree of ancyromonads, phylogenomic analyses based on large multi-marker datasets will be required.

### Species differentiation within the genus *Ancyromonas*

Four new strains and two previously reported strains belonging to the genus *Ancyromonas* were examined in this study (Table 1). *Ancyromonas sigmoides* is the type species of the genus, and the strain B-70 is the neotype strain of this species (Heiss et al., 2010).

Among our new strains, we identified AKRosko2018 as *A. kenti* based on morphology (Figure 1I–P) and molecular phylogeny (Figure 11). We note, incidentally, that strains EDM11b and 3B are actually one and the same: although listed by different strain names in their sequence metadata, both are also given the same culture collection code (CCAP 1953/1) by Glücksman et al. (2013) in their initial description of the species.

By contrast, in spite of their 18S rRNA sequence similarity (98.55% identity), *A. indica* and isolate C362 are quite different in morphology. The posterior flagellum of C362 is as short as or shorter than that of *A. indica*, but unlike *A. indica*, the posterior flagellum is usually visibly acronematic. *Ancyromonas indica* (length/width ratio: ~1.3) is more rounded than C362 (length/width ratio: 1.7). Thus, based on morphology and molecular phylogeny, strain C362 should be treated as a new species of *Ancyromonas* (see Taxonomic Summary), which we propose to call *Ancyromonas mediterranea*.

Finally, our 18S rRNA gene phylogeny suggests that strain SRT303 is likely the sister lineage to the clade containing *A. indica* and *A. mediterranea* C362. The motility of this strain is completely different from any other ancyromonad under our standard culturing conditions. SRT303 can be easily distinguished by the slow movement of the cells (Movie S6). However, as noted above, its motility is no different from that of other ancyromonads when it is cultured in a different medium. This does not diminish the importance of its different behaviour under our standard culturing protocol; rather, it likely reflects a physiological or developmental sensitivity to culturing conditions that is not present in other members of its genus. Such a sensitivity is no less a relevant character than would be a slower movement under all culturing conditions. In addition, while the cell size of SRT303 is within the size range of *A. kenti* and *A. mediterranea*, this strain displayed a longer posterior flagellum. The molecular phylogenetic position and combination of morphological and behavioral traits is novel in this genus, and so we propose this strain to represent a novel species, *Ancyromonas pacifica* (see Taxonomic Summary).

### Proposal of the new genus *Caraotamonas*

In our 50-taxon ML tree (Figure 11), our strain Cro19S5 branched as sister to the *Ancyromonas* clade with maximal statistical support. Its morphological characteristics are, however, different from *Ancyromonas* species. Cells of Cro19S5 possess a much longer posterior flagellum (the longest among all ancyromonad species except for *Fabomonas tropica* NYK3C) and are rounder than any presently known species of *Ancyromonas* (Figure 2I-L). Cells’ movement on the substrate was straighter than that of other ancyromonads (Movie S7), and as such, their posterior flagella typically appeared straight under the light microscope. Thus, based on morphology, behavioral traits, and molecular phylogeny, we regard strain Cro19S5 as representative of a genus distinct from other marine ancyromonads: *Caraotamonas croatica* gen. et sp. nov.

### Species from cultured freshwater lineages

In our phylogenies, three novel strains (STRT, OrsLand19S1 and OrsLand19S2) robustly formed a clade with *Nutomonas.* However, within that clade, the three novel strains “interrupt” *Nutamonas*, separating *Nu. howeae* and *Nu. limna* from the shorter-branching *St. longa*, and with maximal support. The cell size and the length of the posterior flagellum of the new strains each are within the range of other freshwater species. However, they are distinguishable from each other and from the more-closely related *Nutomonas* species (*Nu. howeae* and *Nu. limna*) by their cell shapes (in particular by the flat apex of the rostrum and more pronounced angle between the anterior and dorsal surfaces of the cell) and by differences in their 18S rRNA gene sequences (Table S1). Indeed, the sequences of these strains are considerably divergent, as evidenced by their long branches in our phylogenies. (Figures 11 and 12; Figure S1). Even without our new strains to enforce the separation, Cavalier-Smith (in Glücksman et al., 2013) also noted these differences, and placed the latter species in its own subgenus, as *Nutomonas (Striomonas) longa*. In light of these facts, we elevate *Striomonas* to the rank of genus, thereby formalising the separation of *Striomonas longa* stat. nov. from the remaining species of *Nutomonas* and all other cultured freshwater ancyromonads. For the long-branching freshwater strains, we propose new genus and species names: *Olneymonas thurstoni* for strain STRT, *Nyramonas glaciarium* for strain OrsLand19S1, and *Ny. silfraensis* for strain OrsLand19S2.

### Species differentiation within the genus *Planomonas*

Six species of *Planomonas* have been described so far: *P. micra, P. elongata, P. bulbosa*, *P. brevis, P. melba* and *P. cephalopora.* The latter two species were originally described, by LM only, as *Ancyromonas melba* by Patterson and Simpson (1996) and *Bodo cephaloporus* by Larsen and Patterson (1990), respectively. Both were subsequently transferred to the genus *Planomonas* (Cavalier-Smith et al., 2008; Glücksman et al., 2013). DNA sequences from these two species have never been obtained. It is in fact questionable whether they belong to the genus *Planomonas*. *Planomonas melba* has a normal-width anterior flagellum, so do members of the genus *Fabomonas*. Furthermore, unlike cultured species of both *Planomonas* and *Fabomonas*, the long anterior flagellum of *P. melba* was not described as acronematic, and it has a wider and more ventrally positioned groove (Patterson and Simpson, 1996). *Planomonas melba* has also been observed to swim, unlike all other ancyromonads. Given these differences, it may not belong to any currently recognised genus of ancyromonads. Meanwhile, judging from its original description (Larsen and Patterson, 1990), *P. cephalopora* has an entirely acronematic anterior flagellum; as such, Cavalier-Smith (in Glücksman et al., 2013) should have placed it in the genus *Ancyromonas*, since that was the only marine ancyromonad genus known at the time with that characteristic. In the context of our study, we cannot say whether *P. cephalopora* belongs in *Ancyromonas* or *Caraotamonas*, or indeed in any of the other groups of marine ancyromonads identified in our environmental survey. *Planomonas micra*, the type species of the genus, was described by Cavalier-Smith et al. (2008) using LM, TEM, and molecular phylogenetic analyses. The other three species, *P. elongata, P. bulbosa*, and *P. brevis*, were described by Glücksman et al. (2013), using LM and molecular phylogeny. Although “all four isolates [*P. micra, P. elongata, P. bulbosa*, and *P. brevis*] are nearly indistinguishable microscopically” (Glücksman et al. 2013), they were separated as different species based on ITS2 differences. In our study, we found that our three new strains PMRosko2018, EK-PEI and SRT307, as well as *P. micra, P. elongata, P. bulbosa*, and *P. brevis*, have nearly identical 18S rRNA gene sequences (Table S1), although the four species are clearly distinguishable based on their ITS2 sequences. We found that PMRosko2018’s ITS2 sequence is identical to that of *P. micra*. As such, we identify PMRosko2018 as a strain of *P. micra*. Since we did not obtain ITS2 sequences for either EK-PEI or SRT307, we can identify them no further than the genus level, and regard them as *Planomonas* spp. If SRT307’s ITS2 sequence matches that of *P. elongata*, *P. bulbosa*, or *P. brevis*, we could add the cells’ slow movement (which may not have been observed by Glücksman et al. because it is not exhibited under all culturing conditions) as a distinguishing character to the matching species’ description, an important addition given the overall similarity amongst all ancyromonads.

### Species differentiation within the genus *Fabomonas*

Prior to this study, *Fabomonas tropica* was the sole species in this genus. In this work, we investigated the *F. tropica* type strain, NYK3C^1^, as well as two new strains of the genus, A153 and SRT009. The strain A153 can be easily distinguished from *F. tropica* based on cell size and phylogenetic distance. We thus describe A153 as a new species, *Fabomonas mesopelagica*. However, phylogenetic analyses of the 18S rRNA gene combined *Fabomonas tropica* and SRT009 in an unresolved clade with no internal branch lengths, effectively finding no difference between the strains after trimming. Since we lack the ITS2 sequence for SRT009, we cannot determine its place in the genus on any molecular-genetic basis at this time. In addition, at least under some culturing conditions, SRT009 has a cell movement distinct from that of *F. tropica* NYK3C under the same conditions, and so we consider it premature to name this strain formally, either as *F. tropica* or as a new species, and instead regard SRT009 as an undescribed *Fabomonas* sp., in order to allow further studies to resolve its taxonomic status. A similar situation also applies to two of the three strains included in *F. tropica* by Glücksman et al. (2013), il19 and mex1 (the latter unfortunately dead): each has a distinct morphology, but no significant 18S rRNA gene or ITS2 sequence differences from NYK4. Accordingly, we regard these other two strains likewise as unresolved members of the genus *Fabomonas*.

At least three rows of extrusomes were observed in *F. mesopelagica* in our TEM study (Figure 8B). It is worth noting that only two rows of extrusomes were described in *A. sigmoides* by TEM (Heiss et al., 2011) and in *Planomonas* species by LM (Cavalier-Smith et al., 2008). Similarly, our ultrastructural study of *C. croatica* and *Ny. silfraensis* also demonstrated only two rows of extrusomes. One TEM image of *F. tropica* was published by Glücksman et al. (2013), but its plane of section does not include extrusomes. The same study also included LM of all three strains of *F. tropica*, none of which had clear resolution of extrusomes, and no mention was made of extrusomes in their description. As such, we cannot say whether having triple rows of extrusomes is a characteristic of *F. mesopelagica* in particular, of all members of the genus *Fabomonas*, or of some but not other members of Ancyromonadida as a whole.

## TAXONOMIC SUMMARY

All taxonomic descriptions in this work were approved by all authors.

**Eukarya: Ancyromonadida**

**Family Ancyromonadidae**

*Ancyromonas mediterranea* sp. nov.

**Description:** Elongated ancyromonad cell with a long posterior flagellum that is not visibly acronematic. Cells are 4.5 µm (3.6-5.5 µm) long and 3.0 µm (2.3-3.7 µm) wide.

**Type strain:** C362.

**Isolator:** Maria Ciobanu.

**Type locality:** Mesopelagic water (250 m deep) in Villefranche Bay, France.

**Etymology:** Refers to the type locality of this species in the Mediterranean Sea.

**Gene sequence:** The full ribosomal RNA operon sequence from *Ancyromonas mediterranea* (strain C362) was deposited in GenBank with accession number MW872728.

*Ancyromonas pacifica* sp. nov.

**Description:** Elongated ancyromonad cell with a long posterior flagellum that is not visibly acronematic. Cells are 3.9 µm (3.3-4.5 µm) long and 2.3 µm (1.7-2.8 µm) wide.

**Type strain:** SRT303.

**Isolator:** Takashi Shiratori.

**Type locality:** Muddy saline pool in mangrove stand near beach on Iriomote island, Okinawa Prefecture, Japan.

**Etymology:** Refers to the type locality of this species in the west Pacific Ocean.

**Gene sequence:** The partial small subunit ribosomal RNA gene sequence from *Ancyromonas pacifica* (strain SRT303) was deposited in GenBank with accession number OP740701.

*Caraotamonas* gen. nov.

**Description:** Marine ancyromonad with round cell and entirely acronematic anterior flagellum. Axosomes present and distal to point of flagellar emergence.

**Type species:** *Caraotamonas croatica* sp. nov.

**Etymology:** *Caraota-*used to denote beans in Venezuela; Greek suffix *-monas*, ‘one’ or ‘unit’. *Caraotamonas* is a third-declension feminine Latin noun.

*Caraotamonas croatica* sp. nov.

**Description:** Round and large ancyromonad cell. Posterior flagellum acronematic. Cells are 4.5 µm (3.9–5.4 µm) long and 3.3 µm (2.4–4.0 µm) wide. Two rows of extrusomes. Ancyromonad oscillating movement describes consistently narrow angle.

**Type strain:** Cro19S5.

**Isolator:** Luis Javier Galindo.

**Type locality**: Sediment from the Malo jezero small marine lake in Mljet island, Croatia.

**Etymology:** Refers to the type locality of this species in Croatia.

**Gene sequence:** The full ribosomal RNA operon sequence from *Caraotamonas croatica* (strain Cro19S5) was deposited in GenBank with accession number MW872725.

*Striomonas* stat. nov.

**Change in status:** The subgenus *Striomonas* Cavalier-Smith 2013 of genus *Nutomonas* is promoted to full genus status.

**Description:** as for subgenus *Striomonas* in Glücksman et al. 2013.

**Type species:** *Striomonas longa* stat. nov.

*Striomonas longa* stat. nov.

**Description:** As for *Nutomonas longa* in Glücksman et al. (2013).

**Basionym:** *Nutomonas (Striomonas) longa* Cavalier-Smith and Glücksman 2013.

*Nyramonas* gen. nov.

**Description:** Small freshwater ancyromonads with flat-topped rostra. Entirely acronematic anterior flagellum.

**Type species:** *Nyramonas glaciarium* sp. nov.

**Etymology:** *Nyra-Nýra* means ‘kidney’ in Icelandic, the meaning referring to the cell’s shape and the choice of language reflecting the country of origin for both known species; Greek suffix-*monas*, ‘one’ or ‘unit’. *Nyramonas* is a third-declension feminine Latin noun.

*Nyramonas glaciarium* sp. nov.

**Description:** Small and elongated ancyromonad cell. Cells are 4.0 µm (3.5–4.5 µm) long and 2.5 µm (2.0–2.9 µm) wide.

**Type strain:** OrsLand19S1.

**Isolator:** Luis Javier Galindo.

**Type locality**: Sediment sample from melting glacier in Silfra Rift, Iceland.

**Etymology:** Referring to the collection site in Iceland.

**Gene sequence:** The partial small subunit ribosomal RNA gene sequence from *Nyramonas glaciarium* (strain OrsLand19S1) was deposited in GenBank with accession number OP740706.

*Nyramonas silfraensis* sp. nov.

**Description:** Small ancyromonad cell. Cells are 4.2 µm (3.5–4.9 µm) long and 2.9 µm (2.2–3.6 µm) wide.

**Type strain:** OrsLand19S2.

**Isolator:** Luis Javier Galindo.

**Type locality**: Sediment sample from the walls of (~2 m deep) Silfra Rift, Iceland.

**Etymology:** Referring to the collection location of this species in Silfra Rift, Iceland.

**Gene sequence:** The full ribosomal RNA operon sequence from *Nyramonas silfraensis* (strain OrsLand19S2) was deposited in GenBank with accession number MW872729.

*Olneymonas* gen. nov.

**Description:** Large freshwater ancyromonads. Entirely acronematic anterior flagellum. Ancyromonad oscillating movement describes consistently narrow angle. **Type species:** *Olneymonas thurstoni* sp. nov.

**Etymology:** Refers to the location (Olney Pond) from which the type species was sampled. *Olneymonas* is a third-declension feminine Latin noun.

*Olneymonas thurstoni* sp. nov.

**Description:** Large elongated ancyromonad cell. Cells are 4.9 µm (3.6–5.8 µm) long and 3.0 µm (2.3–3.7 µm) wide. Acronematic posterior flagellum. Contractile vacuole at the anterior end.

**Type strain:** STRT.

**Isolator:** Aaron A. Heiss.

**Type locality**: Rainbow trout caught in Olney Pond, Lincoln, Rhode Island, USA

**Etymology:** Named to honour Stephen Thurston, who provided the sample from which the strain was isolated.

**Gene sequence:** The partial small subunit ribosomal RNA gene sequence from *Olneymonas thurstoni* (strain STRT) was deposited in GenBank with accession number OP740702.

### Family Planomonadidae

*Fabomonas mesopelagica* sp. nov.

**Description:** Bean-shaped cell with a long posterior flagellum. Anterior flagellum nearly same length as the cell. Neither flagellum is visibly acronematic. Cells are 3.8 µm (2.9-4.4 µm) long and 2.6 µm (2.0-3.2 µm) wide.

**Type strain:** A153.

**Isolator:** Maria Ciobanu.

**Type locality**: Mesopelagic water column (250 m depth) in Villefranche Bay, France. **Etymology:** Referring to the collection location of this species in the mesopelagic water column.

**Gene sequence:** The full ribosomal RNA operon sequence from *Fabomonas mesopelagica* (strain A153) was deposited in GenBank with accession number MW872724.

## ACKNOWLEDGEMENTS

This project has received funding from the European Research Council (ERC) under the European Union’s Horizon 2020 research and innovation programme (ERC Starting Grant No 803151 to L.E. and ERC Advanced Grants No 322669 and 787904 to P.L.-G. and D.M., respectively). GT was supported by the 2019 BP 00208 Beatriu de Pinos-3 Postdoctoral Program (BP3; 801370). LG was funded by the Horizon 2020 research and innovation programme under the European Marie Skłodowska-Curie Individual Fellowship H2020-MSCA-IF-2020 (grant agreement no. 101022101—FungEye). We are thankful for access to the Imagerie-Gif core facility at the Institute for Integrative Biology of the Cell (Gif-sur-Yvette, France) and the Electron Microscopy Facility (EMF) of the Institut de Biologie Paris-Seine (Paris, France). We thank members of the UNICELL single-cell genomics platform (https://www.deemteam.fr/en/unicell) for help in cell isolation and rRNA gene sequencing. We also thank Kaleigh Lukacs for assistance with culture maintenance.

## AUTHORS CONTRIBUTIONS

N.Y., G.T., A.H: Conceptualization, Methodology, Investigation, Resources, Writing, Visualisation; L.G.: Conceptualization, Methodology, Investigation, Resources; M.C.C., T.S., J.B.: Resources; P.L-G, D.M., K-I.I, E.K.: Conceptualization, Supervision, Funding acquisition. L.E.: Conceptualization, Writing, Supervision, Project administration, Funding acquisition.

## SUPPORTING INFORMATION

*NB. Supplementary figures and tables are available at figshare (10.6084/m9.figshare.22078451). Supplementary movies are available on the DEEM TEAM YouTube channel (Ancryomonads playlist):* https://www.youtube.com/playlist?list=PLLTaR-PJNuQyhCokjfqXuNjCHc4Iwb2ii

**FIGURE S1.** 18S rDNA phylogenetic trees including environmental sequences, corresponding to Figure 12 and Table S2. Maximum likelihood trees inferred from 1,850 nucleotides using A) TIM3eR4, with support values from 1,000 non-parametric bootstrap replicates, and B) GTR+G4 model, with support values from SH-aLRT analysis and 1,000 ultrafast bootstraps, respectively. Boxes in gray indicate clades consisting exclusively of environmental sequences. Blue font indicates sequences obtained from freshwater samples which branch within a clade of otherwise marine samples. Strains analysed in this study are in bold.

**Movie S1.** Video clip of cells of *Ancyromonas sigmoides* strain B-70.

**Movie S2.** Video clip of cells of *Ancyromonas kenti* strain EDM11b.

**Movie S3.** Video clip of cells of *Ancyromonas* sp., strain JF-An-6.

**Movie S4.** Video clip of cells of *Ancyromonas kenti* strain AKRosko2018.

**Movie S5.** Video clip of cells of *Ancyromonas mediterranea* sp. nov., strain C362.

**Movie S6.** Video clip of cells of *Ancyromonas pacifica* sp. nov., strain SRT303.

**Movie S7.** Video clip of cells of *Caraotamonas croatica* gen. et sp. nov., strain Cro19S5.

**Movie S8.** Video clip of cells of *Nutomonas limna* strain OrsayFeb19Ancy.

**Movie S9.** Video clip of cells of *Striomonas longa* stat. nov., strain ncfw.

**Movie S10.** Video clip of cells of *Nyramonas glaciarium* gen. et sp. nov., strain OrsLandS1.

**Movie S11.** Video clip of cells of *Nyramonas silfraensis* gen. et sp. nov., strain OrsLandS2.

**Movie S12.** Video clip of cells of *Olneymonas thurstoni* gen. et sp. nov., strain STRT.

**Movie S13.** Video clip of cells of *Planomonas* sp., strain EK-PEI.

**Movie S14.** Video clip of cells of *Planomonas micra* strain PMRosko2018.

**Movie S15.** Video clip of cells of *Planomonas* sp., strain SRT307.

**Movie S16.** Video clip of cells of *Fabomonas tropica* strain NYK3C.

**Movie S17.** Video clip of cells of *Fabomonas mesopelagica* sp. nov., strain A153.

**Movie S18.** Video clip of cells of *Fabomonas* sp., strain SRT009.

**Table S1**. Percent identity matrix created by Clustal2.1 on EMBL-EBI web server for 18S rRNA gene sequences from the 45 characterised ancyromonad strains corresponding to Figure 11.

**Table S2**. Environmental survey of ancyromonads in public databases PR2, SILVA, and GenBank (last accessed 2020/10/04), five metabarcoding studies, and clones obtained with newly designed specific primers (see Table S3), corresponding to Figure 12.

**Table S3**. a) Specific primers designed for different ancyromonads (positions refer to *Ancyromonas indica* JQ340337). b) Nine ancyromonad clones recovered from six out of 16 environmental DNAs (Torruella et al. 2017), corresponding to Figures 12 and S1 (in bold).

## Footnotes

1 The type strain is actually NYK4; however, according to its isolator, this is the same as NYK3C (Edvard Glücksman, pers. comm. to AAH).

