## Supplementary figures and images for "Molecular and morphological characterisation of four new ancyromonad genera and proposal for an updated taxonomy of the Ancyromonadida"

### Supplementary Figure 1

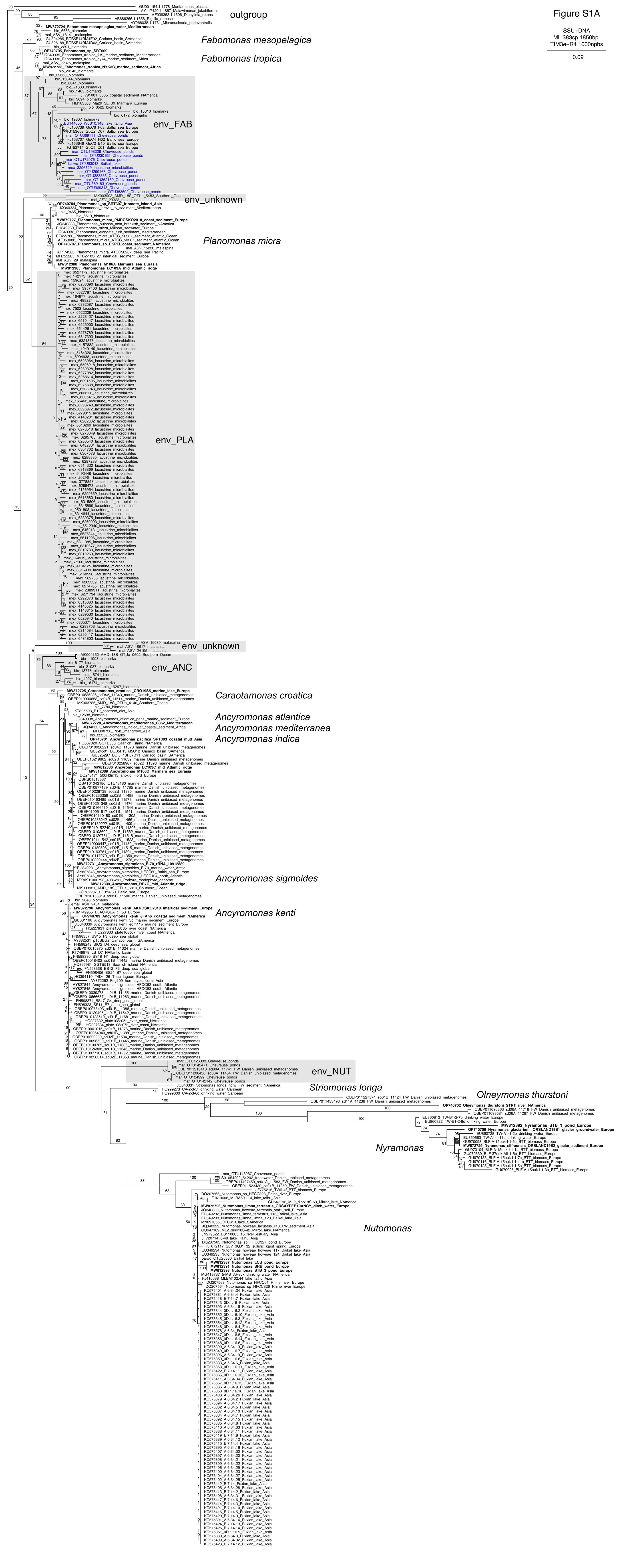

### Supplementary Figure 2

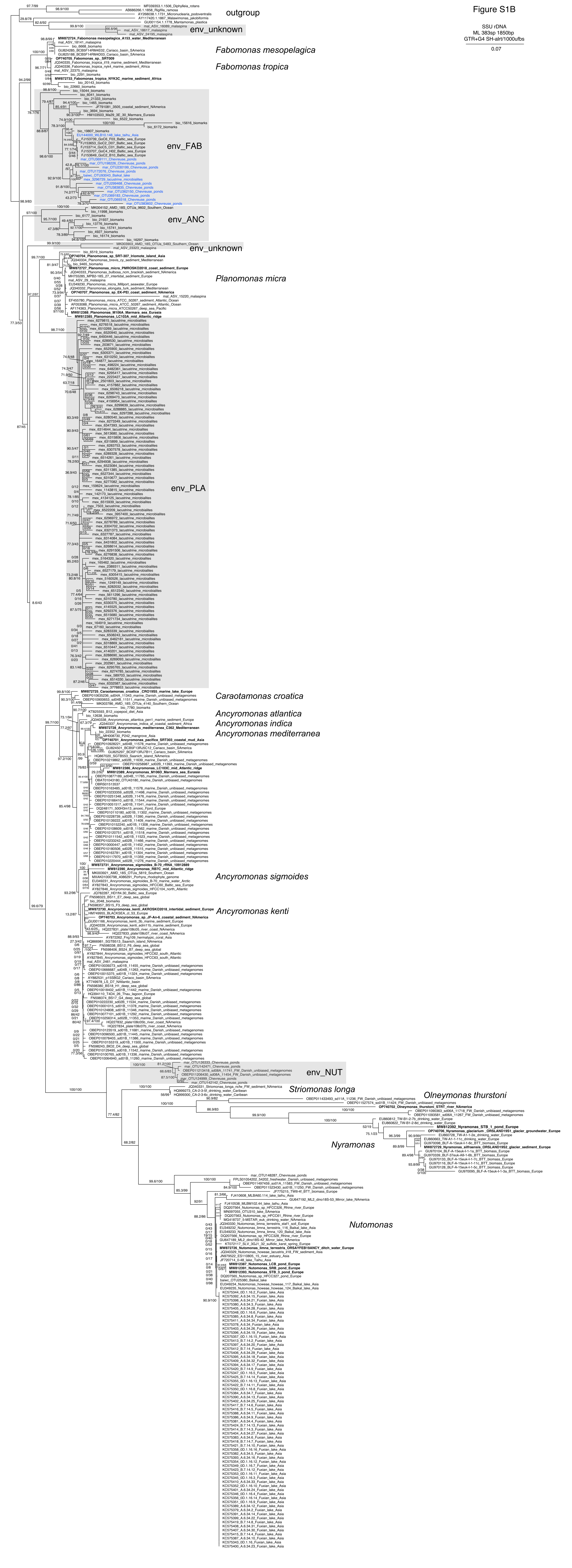
